# Robust partitioning of microRNA targets from downstream regulatory changes

**DOI:** 10.1101/2020.07.23.217117

**Authors:** Ravi K. Patel, Jessica D. West, Ya Jiang, Elizabeth A. Fogarty, Andrew Grimson

## Abstract

The biological impact of microRNAs (miRNAs) is determined by their targets, and robustly identifying direct miRNA targets remains challenging. Existing methods suffer from high falsepositive rates and are unable to effectively differentiate direct miRNA targets from downstream regulatory changes. Here, we present an experimental and computational framework to deconvolute post-transcriptional and transcriptional changes using a combination of RNA-seq and PRO-seq. This novel approach allows us to systematically profile the regulatory impact of a miRNA. We refer to this approach as CARP: <u>C</u>ombined <u>A</u>nalysis of <u>R</u>NA-seq and <u>P</u>RO-seq. We apply CARP to multiple miRNAs and show that it robustly distinguishes direct targets from downstream changes, while greatly reducing false positives. We validate our approach using Argonaute eCLIP-seq and ribosome profiling, demonstrating that CARP defines a comprehensive repertoire of targets. Using this approach, we identify miRNA-specific activity of target sites within the open reading frame. Additionally, we show that CARP facilitates the dissection of complex changes in gene regulatory networks triggered by miRNAs and identification of transcription factors that mediate downstream regulatory changes. Given the robustness of the approach, CARP would be particularly suitable for dissecting miRNA regulatory networks *in vivo*.

## INTRODUCTION

While transcriptional regulation accounts for much of gene regulation, post-transcriptional regulation represents an additional and consequential layer of regulation (1,2). MicroRNAs (miRNAs), a class of small non-coding RNAs, are one of the major *trans*-acting factors responsible for post-transcriptional regulation (3). Together with an Argonaute (AGO) protein, miRNAs function primarily by binding to target mRNA transcripts and inducing mRNA decay and/or translational repression. In humans and other mammals, there are many hundreds of different microRNAs (miRNAs), which collectively regulate the majority of human mRNA transcripts (4) and likely contribute to all gene regulatory pathways. Accordingly, identifying the targets of miRNAs is fundamental to understanding their biological functions, and a wide variety of genomic, biochemical and computational approaches have been developed to address this question (5–7). Despite intense efforts, even the most effective approaches suffer from high rates of false positives and/or negatives (8).

The majority of miRNA target sites in bilaterian animals are found in 3’ untranslated regions (3’UTRs), and comprise a short sequence with perfect complementarity to the 5’ end of the miRNA, or miRNA seed (3). Effective seed-matching target sites are often located within a region of 3 UTR sequence that contains additional features, such as high local AU-content, which determine site efficacy (9,10). In addition to these canonical seed-matching sites, numerous other types of sites have been reported, including sites in coding sequence and 5UTRs, and sites without perfect seed matches (11–14). The extent to which such non-canonical sites contribute to the total targeting repertoire of a miRNA is unclear. Moreover, miRNA-specific parameters influence the targeting properties of certain miRNAs (15,16). The earliest effective approaches to predicting and identifying mammalian miRNA targets used comparative genomics, and worked by cataloguing orthologous 3 UTR sequences whose capacity to basepair perfectly to a miRNA seed sequence is detectably conserved (17–19); such approaches remain an important component of defining biologically consequential miRNA targets. In addition to conserved target sites, a large number of non-conserved sites also respond to their cognate miRNAs (20). Non-conserved sites constitute the majority of total sites; therefore, conservation alone cannot be used to robustly identify target sites (4). Numerous computational approaches exist which predict the strength or efficacy of miRNA target sites, with varying degrees of effectiveness (6,7). In general, these approaches are built on experimental data in which the transcriptome or proteome is monitored in response to high and transient exposure to an exogenous miRNA. The resulting data, aggregated over many experiments, is used to train a model that captures the response to a miRNA, and the results extrapolated to other miRNAs and other cell types. Such tools have played an important role in accelerating our understanding of miRNA biology. Biochemical techniques (including CLIP-based assays) have also been used to identify miRNA targets (21–24); however, these assays suffer from high levels of background, perhaps arising from the transient nature of AGO binding. Indeed, subsequent attempts to verify non-canonical target sites identified from CLIP have shown that such sites are largely ineffective (16).

Although approaches that identify or predict miRNA targets have continued to evolve and improve, the vast majority of both training and validation datasets rely upon cell culture experiments in which an exogenous miRNA is introduced transiently at high concentration (5). Extending these approaches to *in vivo* settings, with miRNA knockouts, for example, has indicated that target prediction remains valuable but imperfect. Deviations between target prediction and *in vivo* miRNA-mediated regulation presumably derive from numerous sources. Biologically consequential miRNAs are enmeshed within complex gene regulatory networks, and the action of such a miRNA is likely to elicit substantial downstream changes beyond the direct targets (25). For example, a miRNA may directly repress an mRNA encoding a transcription factor, thus altering the downstream targets of the transcription factor and potentially confounding efforts to identify the direct targets of the initiating miRNA. Indeed, a large body of literature illustrates intimate mingling of miRNAs and transcription factors within gene regulatory networks (26); thus, identifying transcription factors that direct downstream regulatory changes initiated by a miRNA is likely an important step towards understanding biological functions of miRNAs. This complexity in miRNA regulatory networks alone makes miRNA target prediction *in vivo* problematic. Three major challenges exist: (i) the relatively subtle regulation elicited by a miRNA, often less than 2-fold, (ii) the large number of potential targets, often several hundred, and finally, (iii) for consequential miRNAs, the extent of downstream changes.

A popular and effective approach to identifying miRNA targets *in vivo* is to intersect lists of genes differentially expressed in response to a specific miRNA with lists of predicted targets. Importantly, both lists often include many hundreds of genes; thus, random overlap alone will generate a substantial set of intersecting candidate direct miRNA targets. We reasoned that eliminating genes whose differential expression derives from transcriptional regulation might enable more robust delineation of the direct targets of a miRNA. Prior to our work, a conceptually similar approach has been developed: EISA (Exon–Intron Split Analysis) exploits intron-mapping reads in RNA-seq data to indicate levels of pre-mRNAs and thus serves as a proxy for transcriptional activity (27). The advantage of EISA is that it is straightforward to implement; nevertheless, pre-mRNA levels are not a direct sensor of transcription, potentially compromising the accuracy of this method.

Here, we use PRO-seq (Precision Run-On sequencing), a tool that directly monitors transcription across the genome (28), in combination with RNA-seq to robustly distinguish between direct miRNA targets and indirect effects arising from downstream regulation. We corroborate the efficacy of our approach using orthogonal genomic assays to measure AGO-miRNA binding to targets (AGO eCLIP-seq (29)) and translational efficiency (ribosome profiling (30)). We use these data to investigate mechanisms of miRNA-mediated repression; for example, we quantify the contributions of miRNA-mediated mRNA decay and translational repression using miRNAs expressed at physiological levels. Additionally, we identify novel, effective miRNA target sites residing within the open reading frame (ORF); interestingly, such coding sites are only prevalent for a subset of miRNAs that we examine. Because PRO-seq also profiles activity of DNA regulatory elements, such as enhancers and promoters (31), we identify candidate transcription factors associated with regulatory elements exhibiting altered transcriptional activity. We find that activities of these transcription factors are modulated by specific miRNAs targeting the cognate transcript, and that such repression contributes to the downstream changes in transcriptional regulation. Using CARP to deconvolute regulation occurring at the level of transcription, post-transcription, or both, we demonstrate that the combined analysis of RNA-seq and PRO-seq is a powerful approach to investigate complex transcriptional and post-transcriptional gene regulatory networks.

## MATERIAL AND METHODS

### Cell culture

Flp-In T-REx 293 (HEK293-derived; Invitrogen) and HEK293T (American Type Culture Collection, ATCC) cells were used for all experiments in this study. Cells were cultured at 37°C in a humidified incubator containing 5% CO_2_ and maintained in DMEM (Life Technologies) containing 10% FBS (Sigma-Aldrich) and 1% penicillin/streptomycin (Life Technologies); Flp-In T-REx 293 cells were also supplemented with 100 μg/ml Zeocin (Invitrogen). Cells were passaged every 2-4 days. Cell lines were not tested for mycoplasma contamination.

### Synthetic miRNA hairpin constructs

Expression cassettes consisting of the doxycycline-inducible cytomegalovirus (CMV) promoter (derived from the Invitrogen T-REx system), the intron-containing EF1α 5 UTR, and *Aequorea coerulescens* GFP were cloned into a lentiviral transfer vector containing a neomycin selective resistance cassette. To clone synthetic miRNA hairpins into the intron, *Xba*I and *Xma*I restriction sites were introduced. An artificial hairpin backbone (“A5” from (32) was used and the sequence of the mature miRNA was replaced with the sequences of the seven miRNAs used in this study: hsa-miR-1, hsa-miR-122, hsa-miR-133a, hsa-miR-155, hsa-miR-302a, hsa-miR-372 and hsa-miR-373 (hairpin sequences included in Supplementary Table S1). To insert the hairpin sequences, pairs of complementary oligonucleotides were designed containing the sequences with terminal restriction enzyme sites *Xba*I and *Xma*I. The oligos were annealed, extended, digested and ligated into the GFP vector. The parent GFP vector lacking a miRNA hairpin was used as a negative control for all experiments.

### Generation of stable cell lines expressing specific miRNAs

Lentiviruses were generated by transfecting the lentiviral transfer vectors described above along with packaging (psPAX2, Addgene plasmid #12260) and envelope (pMD2.G, Addgene plasmid #12259) plasmids with Lipofectamine 2000 (Invitrogen) in HEK293T cells. The media was replaced 24 hours later with fresh DMEM containing 30% FBS, and lentiviral supernatants were collected 24 hours later. To generate stable cell lines expressing miRNA constructs, Flp-In T-REx 293 cells were transduced and selected with 1 mg/mL Geneticin (Invitrogen) for six days, after which the Geneticin concentration was lowered to 0.8 mg/ml. To ensure steady-state miRNA levels, miRNA-GFP cassettes were induced by adding 1 μg/mL doxycycline every day for seven days.

Biological replicates were transduced and maintained separately for all experiments. Two biological replicates were generated for experiments employing miR-1 and miR-122, while three replicates were generated for miR-133, miR-155, miR-302a, miR-372 and miR-373 experiments.

### miRNA qPCR

Primers were designed as described previously (33) (primer sequences in Supplementary Table S2). The reverse transcription primer for all miRNAs is 5’-CAGGTCCAGTTTTTTTTTTTTTTTVN, where V is A, C and G and N is A, C, G and T. The reverse primer for all miRNAs is 5’-CAGGTCCAGTTTTTTTTTTTTTTT. For cDNA synthesis, 100 ng of total RNA from all samples and 100 amol spike-in RNA (sequence in Supplementary Table S2) were heat denatured at 65°C for 30s. The reaction mix included 1 μl of 10x poly(A) polymerase buffer (NEB), 0.1 mM of ATP, 1 μM of RT primer, 0.1 mM of each deoxynucleotide (dATP, dCTP, dGTP and dTTP), 100 U of RevertAid Reverse Transcriptase (Thermo Scientific) and 1 unit of poly(A) polymerase (New England Biolabs) and was incubated at 42°C for 1 hour followed by enzyme inactivation at 95°C for 5 minutes. For quantitative PCR, synthetic templates used for the standard curve were DNA oligonucleotides complementary to the miRNAs with a reverse primer binding site incorporated into the 5’ end (sequences in Supplementary Table S2). Synthetic templates were used to generate a standard curve, with dilutions ranging from 0.5 fM to 50 pM final concentrations with 0.2 ng salmon sperm DNA. Quantitative PCR of biological replicates was performed in 10 μl total volume with 0.1 μl of cDNA. Cycling conditions were 95°C for 5 min followed by 40 cycles of 95°C for 10 sec and 60°C 30 sec and 70°C 30 sec. A melting curve analysis (55°C to 95°C) was performed after the thermal cycling. Quantitative PCR was performed on a LightCycler 480 Instrument II (Roche).

### PRO-seq

#### Library preparation

Biological duplicates of cells expressing miR-1, miR-122 and empty vector control were harvested after seven days of induction with doxycycline. For each sample, cells were scraped from one 10 cm plate in ice-cold PBS. A portion of the cells (20%) was set aside for RNA-seq and the remaining 80% was used for PRO-seq following a protocol adapted from (34). Briefly, nuclei were isolated using a buffer containing 0.05% tween-20, and incubated with biotin-labelled nucleotides in a nuclear run-on reaction along with sarkosyl. The total RNA was extracted using Trizol and fragmented using NaOH hydrolysis for 20 minutes on ice. The biotin-labelled fragments of nascent RNAs were enriched using Streptavidin M280 beads (Invitrogen) followed by ligation of 3’ ends with pre-adenylated DNA adapters (App-GATCGTCGGACTGTAGAACTCTGAAC/3InvdT/) using T4 RNA Ligase 2 truncated K227Q (NEB) in absence of ATP. Following another round of biotin-enrichment, the 5’ ends of the RNA were modified and ligated with 5’ RNA adapter (CCUUGGCACCCGAGAAUUCCA). The cDNA was generated using SuperScript III RTase (Invitrogen). The libraries were PCR amplified, size selected using PAGE and sequenced on an Illumina NextSeq 500.

For miR-133a, miR-155, miR-302a, miR-372 and miR-373, libraries were prepared in biological triplicate (along with the empty vector control), as described above, except for the following modifications, adapted from (35). A 3’ RNA adapter (pNNNNNNXXXXXXNNGAUCGUCGGACUGUAGAACUCUGAAC/3InvdT/) containing sample barcodes (“X”s) was ligated to the RNA 3’ termini using T4 RNA ligase I (Invitrogen), allowing us to pool samples post ligation, streamlining subsequent steps in the protocol. We used unique molecular indexes (UMIs) in both the 3’ and 5’ RNA adapters (5’ adapter: CCUUGGCACCCGAGAAUUCCANNNNN) to minimize ligation bias and facilitate removal of PCR duplicates from the sequencing data.

#### Data processing

For miR-1, miR-122 and the empty vector control, single-end sequencing reads were trimmed to remove adapter sequences using fastx_clipper tool from FASTX Toolkit v0.0.13 (-a TGGAATTCTCGGGTGCCAAGG -l 0 -Q33) for miR-1 and miR-122, and cutadapt v1.12 (-a TGGAATTCTCGGGTGCCAAGG -m 0 -f fastq) for miR-133a, miR-155, miR-302a, miR-372 and miR-373. All trimmed sequences shorter than 15 nucleotides were removed and the remaining reads were reverse complemented. The reads originating from PhiX or rRNAs were removed using Bowtie v1.1.2-based alignments (36) to respective reference sequences. The resulting high-quality reads were aligned to hg19 genome using Bowtie v1.1.2 (-p7 -v2 -m1 -q), and the resultant BAM files were used for further analysis.

For miR-133a, miR-155, miR-302a, miR-372 and miR-373 and the empty vector control, reads were first demultiplexed using the in-line sample barcode. The UMI sequences were removed from each read and placed into the read name. Subsequent processing steps were as described for miR-1 and miR-122 samples, with an additional removal of PCR duplicates, defined as reads containing identical UMI sequences and mapping locations.

#### Visualization of PRO-seq data on genome browser

The PRO-seq BAM alignments were converted to BED format using bamToBed (BEDTools v2.26.0; (37)) and were split into two files based on the genomic strand they belong to. The read alignments were converted to genome coverage using genomeCoverageBed (-i stdin -3 -bg; BEDTools v.2.26.0). The resulting bedgraph files were normalized by read depth and were converted to bigwig files using bedGraphToBigWig (38) for visualization on the UCSC genome browser.

#### Peak calling using dREG

PRO-seq BAM alignments were converted to BED format using bamToBed (BEDTools v.2.26.0) and were split into two files based on the genomic strand they belong to. The read alignments were converted to genome coverage using genomeCoverageBed (-i stdin -3 -bg; BEDTools v.2.26.0). The resulting bedgraph files were converted to bigwig files using bedGraphToBigWig (38), without read depth normalization. dREG (v18.11.2016; (39) was used to call peaks using the bigwig files and default dREG parameters. Transcriptional activity was estimated by counting the number of reads mapping to the dREG peaks using featureCounts (-F SAF -s 0 -Q 50 -T 10) of Subread package v1.5.1 (40). Peak centers were identified using dREG (v6.1.2018; (41)) and default parameters.

#### Quantification of transcriptional output

To calculate transcriptional output from PRO-seq, we considered only gene-body reads, removing all reads originating from regulatory elements such as promoter or enhancer elements. These transcriptional regulatory elements were predicted using dREG. BEDTools was used to remove reads mapping to these regulatory elements, and the remaining gene-body reads were counted using featureCounts (-F SAF -s 1 -Q 50).

### RNA-seq

#### Library preparation

Cells scraped from 10 cm plates in PBS were pelleted and resuspended in 1 ml Trizol (Invitrogen). The total RNA was extracted from Trizol according to manufacturer’s instructions, with the addition of a chloroform extraction. Directional RNA-seq libraries were prepared from 1000 ng total RNA per sample using the NEBNext Ultra II Directional RNA Library Prep Kit for Illumina (New England Biolabs), with initial polyA+ isolation, by the Transcriptional Regulation and Expression Facility at Cornell University.

#### Data processing

Raw reads were trimmed to remove adapter sequences using fastx_clipper (parameters: -l 15 -Q33 -a GATCGGAAGAGCACACGTCTGAACTCCAGTC) and were aligned to the human genome (hg19) using TopHat v2.1.1 (--library-type fr-firststrand; (42)); the resultant BAM files were used for the further analyses. The featureCounts software was used to count reads mapping to exons (parameters: -F SAF -s 2 -Q 50) and introns (parameters: -F SAF - s 2 -Q 50 --fracOverlap 1). The differential expression analysis was performed using edgeR v3.24.3 (43) and q-values were computed using qvalue v2.14.1 R package (44).

#### Predicting 3’UTR isoforms

The BAM files across all RNA-seq samples were merged and the read alignments with mapping quality ≥ 50 were extracted using samtools v1.9 (45). The resultant read alignments were used to predict the poly(A) cleavage sites using GETUTR v2.0.0 (46). The predicted poly(A) cleavage site with the highest score (representing the major 3 UTR isoform) was used to modify 3 UTR annotations (gencode v19). Specifically, annotated 3UTRs were trimmed up to the genomic locations of the highest-scored poly(A) cleavage site. Annotations were not adjusted for inferred poly(A) cleavage sites downstream of annotated 3UTRs. The modified annotations were used to exclude genes that loose predicted miRNA target sites due to alternative polyadenylation and cleavage.

### Statistical significance of post-transcriptional regulation

Transcriptional output estimated using PRO-seq and mRNA abundance quantified using RNA-seq were used for determining genes that exhibit significant post-transcriptional regulation. Lowly expressed genes (counts per million ≤ 1 in any sample) were filtered out and the remaining genes were used for a statistical test using edgeR, based on code provided in (27). The edgeR analysis was performed under generalized linear modeling framework. Like typical edgeR expression analysis, one factor was defined based on the experimental conditions, and the second factor represented type of assays (RNA-seq versus PRO-seq). A model containing these two factors and an interaction term between the two (full model) was compared with the reduced model, the full model without the interaction term, using likelihood ratio test (LRT). The q-values were computed using qvalue R package (44).

### Exon-intron split analysis (EISA)

The EISA was performed by using exonic read counts and intronic read counts obtained from RNA-seq experiments (see above). The code provided in (27) was used for this analysis.

### small RNA sequencing

#### Library preparation

Cells were lysed in confluent 6-well plates using 500 uL Trizol (Invitrogen). Total RNA was extracted from Trizol according to manufacturer’s instructions, with the addition of a chloroform extraction. Libraries were prepared from 1000 ng total RNA using the NEBNext Multiplex Small RNA Library Prep Set for Illumina (New England Biolabs) by the Transcriptional Regulation and Expression Facility at Cornell University.

#### Data processing

The small RNA data were processed using miRDeep2 (47). Briefly, the adapters were trimmed and duplicate reads were collapsed using mapper.pl (-d -e -h -i -j -k AGATCGGAAGAGCACACGTCT -l 18 -m -s reads.fa -v). The collapsed reads were aligned to miRNA hairpins and reads originating from mature miRNAs were counted using quantify.pl (-t hsa -d -W). The miRNA hairpins and mature miRNA sequences used in quantify.pl were downloaded from miRbase v21 (48). The sequences of synthetic hairpins used in this study were appended to the miRNA hairpin file. The miRNA processing efficiency was determined by calculating the fraction of read ends mapping at each position of the hairpin using the miRBase.mrd file generated by quantify.pl.

### AGO eCLIP

#### Library preparation

Biological duplicates of cells stably integrated with miR-1, miR-122, or the empty vector control were passaged in the presence of 1 μg/ml doxycycline for seven (replicate 1) or eight (replicate 2) days. For each sample, two ~70% confluent 10 cm plates were UV crosslinked on ice at 400 mJ/cm^2^, washed with PBS, lifted from the plates, pooled, pelleted, snap frozen in liquid nitrogen and stored at −80°C. For preparing eCLIP libraries, cell pellets were thawed, lysed and prepared with a protocol adapted from (29). Briefly, cell pellets were thawed and lysed in 1 ml lysis buffer (50 mM Tris-HCl pH 7.4, 100 mM NaCl, 0.5% Igepal CA-630) supplemented with protease inhibitor cocktail III (EMD Millipore), treated with RNase I (Ambion), Turbo DNase (Invitrogen), and clarified. At all points, RiboLock RNase inhibitor (Thermo Scientific) was used instead of Murine RNase Inhibitor. For immunoprecipitation, 10 μg Ago2 antibody (Anti-AGO2 clone 11A9, MABE253, EMD Millipore) was bound to 100 μL washed Dynabeads Protein G (Invitrogen). The clarified lysate (950μl) was added to washed antibody-coupled beads and rotated at 4° C for 4 hours. The IP was washed 2x with high salt wash buffer (50 mM Tris-HCl pH 7.4, 1 M NaCl, 1 mM EDTA, 0.5% Igepal CA-630), 1x wash buffer (20 mM Tris-HCl pH 7.4, 10 mM MgCl2, 0.2% Tween-20), and then washed with 1x FastAP buffer (10 mM Tris pH 7.5, 5 mM MgCl_2_, 100 mM KCl, 0.02% Triton X-100). Beads were treated with FastAP (Thermo Scientific), Turbo DNase, and T4 PNK (NEB). A 3’ RNA adapter (5’ P-TGGAATTCTCGGGTGCCAAGG/3InvdT) was ligated to the RNA on-bead, and resuspended in LDS sample buffer (Life Technologies). Inputs and 10% IP were run on SDS-PAGE, transferred to nitrocellulose membrane, and visualized by western blot to verify pulldown. For preparing eCLIP libraries, 90% of the IP was run on SDS-PAGE and transferred to nitrocellulose membrane. For each sample, the membrane was cut from ~97kDa up to ~275kDa to isolate AGO2-RNA complexes. Membrane slices were treated with proteinase K and urea, extracted with acid phenol:chloroform and cleaned as described (29). RNA was reverse transcribed with SuperScript III (Invitrogen) with the RT primer 5’ CCTTGGCACCCGAGAATTCCA. cDNA was cleaned up and a 3’ DNA adapter (5’ P-NNNNNNNNNNGATCGTCGGACTGTAGAACTCTGAAC/3InvdT) containing a 10 nucleotide unique molecular identifier (UMI) was ligated to the 3’ end of the cDNA, and cleaned as described (29). Approximately 90% of the cDNA was amplified for 16 cycles using sample-specific Illumina-compatible primers, ethanol precipitated and run on an 8% PAGE gel. The library smear from ~160bp to ~400bp was cut from the gel, purified, quantified, pooled, and sequenced on the Illumina NextSeq500 with the 75bp kit.

#### Data processing

Following removal of UMI sequences from the read sequence, the reads were subjected to adapter trimming using cutadapt v1.8.3 (-a TGGAATTCTCGGGTGCCAAGG -m 18). All reads mapping to ribosomal RNAs using Bowtie2-based alignment (v2.3.5.1; (49)) were removed and the remaining reads were aligned to the human genome (hg19) using TopHat v2.1.1 (-g 1 -p 2 --library-type fr-secondstrand). The alignments were processed using samtools. PCR duplicates, as defined by reads containing identical UMI barcodes and mapping locations, were removed using an in-house Perl script. Peaks were called with CLIPper (50), and reproducible peaks (IDR < 0.1) were obtained using Irreproducibility discovery rate (IDR) analysis (51). The reproducible peaks from all samples were pooled and overlapping peaks were merged using BEDTools to produce a set of consolidated eCLIP peaks, which was used for the further analyses. Relative AGO density was calculated for each peak by computing a ratio of normalized read counts in miRNA-expressing cells to those in control cells. Because increased AGO binding results in reduced mRNA levels due to miRNA repression, this measurement of AGO density is underestimated.

### Ribosome profiling

#### Library preparation

Ribosome profiling libraries were prepared with the TruSeq Ribo Profile (Mammalian) Kit (Illumina). RNA-seq libraries for normalizing ribosomal footprints were prepared in parallel according to kit instructions. Biological duplicates of cells stably integrated with miR-1, miR-122, or the control were induced with 1 μg/ml doxycycline for seven days. To stall ribosomes, cells in 10 cm plates were incubated in media supplemented with 100μg/ml cycloheximide for two minutes at 37°C. Cells were washed in ice-cold PBS supplemented with 100 μg/ml cycloheximide, lifted from the plates, pelleted, and lysed in 750 μL mammalian lysis buffer on ice for 10 min. Lysate was clarified and split into two tubes: (i) 100 μL for preparing total RNA libraries, and (ii) 200 μL for prepare ribosome footprint libraries. Library preparation was performed according to the protocol. Ribosomes were treated with 60 U TruSeq Ribo Profile Nuclease to generate ribosome-protected fragments and isolated via size-exclusion with an Illustra MicroSpin S-400 HR column (GE healthcare). Ribosomal RNA was depleted from both ribosome-protected fragments and RNA-seq libraries using the Illumina Ribo-Zero Gold Kit (Human/Mouse/Rat) according to the protocol. Libraries were sequenced on the Illumina NextSeq 500 with the 75bp kit.

#### Data processing

The raw reads were trimmed to remove adapter sequences using cutadapt v1.8.3 (-a AGATCGGAAGAGCACACGTC -m 18). Trimmed reads originating from rRNA were removed using Bowtie2 v2.3.5.1 with default parameters for RNA-seq datasets and with L 20” and other default parameters for Ribo-seq datasets. Remaining reads were mapped to the hg19 genome and gencode v19 annotated genes using Tophat v.2.1.1 (--no-novel-juncs -transcriptome-index <indexFile> -p 3 --library-type fr-firststrand). Reads mapping to coding region excluding the ends (initial 45nt and ending 15nt) were counted using featureCounts (-F SAF -s 1 -Q 50 -T 10 --fracOverlap 1). The change in translational efficiency was calculated using edgeR in the same manner as the calculation for change in post-transcriptional regulation. Briefly, the first factor represented experimental conditions and the second factor represented the type of assay (ribosome profiling verses RNA-seq). The models with or without the interaction term between the two factors were compared using likelihood ratio test in edgeR framework. The q-values were computed using qvalue R package (44).

### Tissue-specificity analysis of miRNA direct targets

This analysis was performed as described previously (52). Briefly, for each gene, 53 different tissues were ranked based on the expression of that gene in the tissue expression data obtained from the GTEx Portal (downloaded on Feb/2020). Using these ranks as expression levels, for each tissue, the distribution of ranks of miRNA direct targets were compared with that of 5000 randomly selected genes. To test if the expression levels of miRNA direct targets are significantly lower compared to that of the randomly selected genes, one-sided Wilcoxon rank sum tests were performed, and the resultant P values were corrected using the Benjamini-Hochberg method. The genes with zero counts in more than 26 tissues were excluded from the analysis.

### Prediction of 5’UTR and ORF sites

Since TargetScan predictions do not include 5 UTR and ORF sites, the predictions for sites located in 5 UTR and ORF were performed based on matches with 8mer, 7mer-m8 and 7mer-A1 sites motifs. The conserved ORF sites were obtained from PACCMIT-CDS (predictions_human_cons.txt) (53) by filtering predictions with P_SH values larger than 0.05.

### Motif enrichment analysis for transcription factor binding sites

The motif enrichment analysis was performed as described previously (54) with modifications. Putative transcription factor binding sites (TFBS) were identified in dREG peaks by first obtaining the 3059 motifs corresponding to binding sites for 1735 distinct transcription factors (TF) CISBP (55). These TF binding motifs were then searched in a window of 150 bp centered on dREG peak centers to identify putative TFBS, using FIMO with p-value cut-off of 10e5 (meme_4.12.0; (56)). TFBS enrichment was computed over different subsets of peaks. Subsets of peaks, denoted by S, that are significantly (FDR < 0.05) more/less active in miRNA-expressing cells compared to the control cells were identified using edgeR. Computations were performed to calculate the fraction of peaks in a subset s, and fraction of peaks in a set of 10,000 randomly selected peaks that contain at least one binding motif for a TF t, denoted by fs and fr, respectively. The ratio of fs/fr was used to determine enrichment (ratio > 1) or depletion (ratio < 1) of binding motif for TF t in a subset s. The significance P value of enrichment or depletion was computed using a binomial test, where the set of 10,000 randomly selected peaks was used as the null distribution, followed by FDR correction. Transcription factors depicted in Figure 6C were direct targets of respective miRNAs (CARP q-value < 0.05 and presence of predicted target sites) and whose binding motif was significantly enriched or depleted (FDR < 10^-3^) in the given subset, s. The heatmaps in Figure 6C were prepared using an R package, ComplexHeatmap (57).

### Comparing two distributions

Statistical significance for the difference between two distributions were calculated using two-sided Wilcoxon rank sum test (wilcox.test function in R) for all comparisons unless mentioned otherwise.

### TargetScan

The TargetScan predictions were downloaded from http://www.targetscan.org/vert_70/. Predictions with Pct value of “NULL” were excluded.

### Luciferase reporter assays

For verifying miRNA activity of stably-integrated miRNAs (Supplementary Figure S1F), a sequence containing two miR-1 target sites was excised from pAG76 (20) using restriction enzymes *Sac*I and *Xba*I and inserted into pmirGLO Dual-Luciferase vector (Promega) using restriction enzyme sites for *Sac*I and *Xba*I (sequence for miR-1 sites in Supplementary Table S3). The miR-1 sites were disrupted to generate a negative control (sequence in Supplementary Table S3). The constructs used in Supplementary Supplementary Figure S1G contained a perfectly complementary site inserted in pmirGLO vector at *Xba*I and *Sal*I restriction sites (sequences in Supplementary Table S4).

For assaying miR-122 ORF target sites, candidate sites were chosen from those genes that contain exactly one predicted target site in the ORF and none in the 3 UTR, and that exhibited post-transcriptional repression. A 78nt region (sequences in Supplementary Table S5) centered on the candidate miR-122 ORF site was cloned with NEB HiFi DNA Assembly Master Mix (New England Biolabs) into the open reading frame of firefly luciferase in pmirGLO. A short linker (amino acid sequence GGGSGGGS) was added to firefly luciferase after the last amino acid, followed by the 26 amino acid sequence taken from the ORF site, followed by a stop codon. As a control, 2-4 nucleotide synonymous mutations were introduced within the miRNA seed sequence to disrupt the site, with attempts to maintain similar codon usage frequencies.

For all reporter assays, 1×10^5^ HEK293 cells were seeded per well in 24-well plate 24 h prior to transfection. For experiments in Supplementary Figure S1F and G, HEK293 cells expressing specific miRNAs were transfected with 30 ng of pmirGlo reporter plasmids using 0.5 μL of Lipofectamine 2000 (Invitrogen) and were harvested 24 h after transfection. For the experiments in Figure 4 and Supplementary Figure S4, cells were transfected with 100 nM miR-122 mimics or a non-specific miRNA mimic (sequences in Supplementary Table S5) and 6 ng pmirGlo reporter plasmid and harvested 24 h after transfection. Assays were performed using the dual-luciferase reporter assay kit (Promega) and a Veritas Microplate Luminometer (Turner Biosystems). Firefly luciferase values were normalized to *Renilla* luciferase as a transfection control. For Figure 4C and Supplementary Figure S4F, luciferase values for cells transfected with the wild-type or mutant reporters and the miR-122 mimic were also normalized to the geometric mean of cells co-transfected with the identical reporter plasmids and the non-specific miRNA mimic.

### Modeling miRNA expression over time

The empirical determination of the time required for miRNA levels to reach steady-state was performed by combining the synthesis (β) and decay (α) rates of miRNAs in the following exponential function.

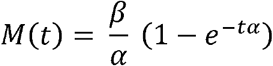

Where, M(t) represents the level of miRNAs at time t (hours). The synthesis rate β was assumed to be 1 unit and the decay rate α was calculated using ln(2)/h, where h represents half-life of a miRNA in hours. Different half-lives ranging from 6 to 36 hours, comprising the majority of miRNAs (58) were used to calculate levels of miRNAs over 1 to 8 days period. The results are plotted in Supplementary Figure S1A.

## RESULTS

### A system for measuring miRNA-mediated post-transcriptional regulation

Given the limitations of conventional methods for identifying direct miRNA targets, we sought to develop a novel experimental approach to robustly identify genes subject to post-transcriptional regulation. While direct miRNA targets are regulated exclusively at the post-transcriptional level, we began this work under the assumption that indirect targets would be predominantly regulated at the transcriptional level. We used RNA-seq and PRO-seq to measure steady-state mRNA levels and transcriptional output, respectively, and integrated these datasets to deconvolute the post-transcriptional regulation (Figure 1A). To develop this method and to evaluate its efficacy, we profiled HEK293 cells in the presence and absence of specific miRNAs. We chose first to study miR-1 and miR-122 because they are well-studied human miRNAs not expressed in HEK293 cells. The majority of datasets used to identify miRNA targets and to train prediction algorithms are generated using cell culture with transiently transfected miRNA, which are typically introduced at high levels. We elected to stably integrate miRNA hairpins embedded within the intron of a doxycycline-inducible GFP reporter, to more closely approximate *in vivo* expression levels. To ensure accurate processing of the mature miRNAs, we designed miRNA hairpins based on established sequence and structural features favored by the miRNA biogenesis machinery (32). In order to approximate steady-state miRNA levels (58), we treated cells with doxycycline for seven days (Supplementary Figure S1A). We confirmed high physiological levels and accurate processing of induced miRNAs using small RNA sequencing (small RNA-seq; Supplementary Figure S1B and C). Because accurate quantification of miRNA levels by small RNA-seq is compromised by ligation biases during library preparation (59), we also used quantitative PCR (qPCR) to establish that miR-1 and miR-122 are expressed within the physiological range of other miRNAs found in HEK293 cells (Supplementary Figure S1D): compared to miR-10, the most highly detected endogenous miRNA in our small RNA-seq data, miR-122 is expressed at about the same level and miR-1 is two-fold higher. Furthermore, using small RNA-seq and RNA-seq, we noted that the overall miRNA profile and activity of endogenous miRNAs (Supplementary Figure S1E), respectively, were unchanged upon induction of these exogenous miRNAs. Importantly, using reporter assays, we found that the induced miRNAs are functionally active (Supplementary Figure S1F) and that the anticipated miRNA guide strand was chosen selectively for loading onto AGO (Supplementary Figure S1G). Collectively, these results indicate that our synthetic miRNA expression system mimics endogenous miRNA, facilitating the analysis of miRNA targets while approximating normal *in vivo* parameters.

**Figure 1.**
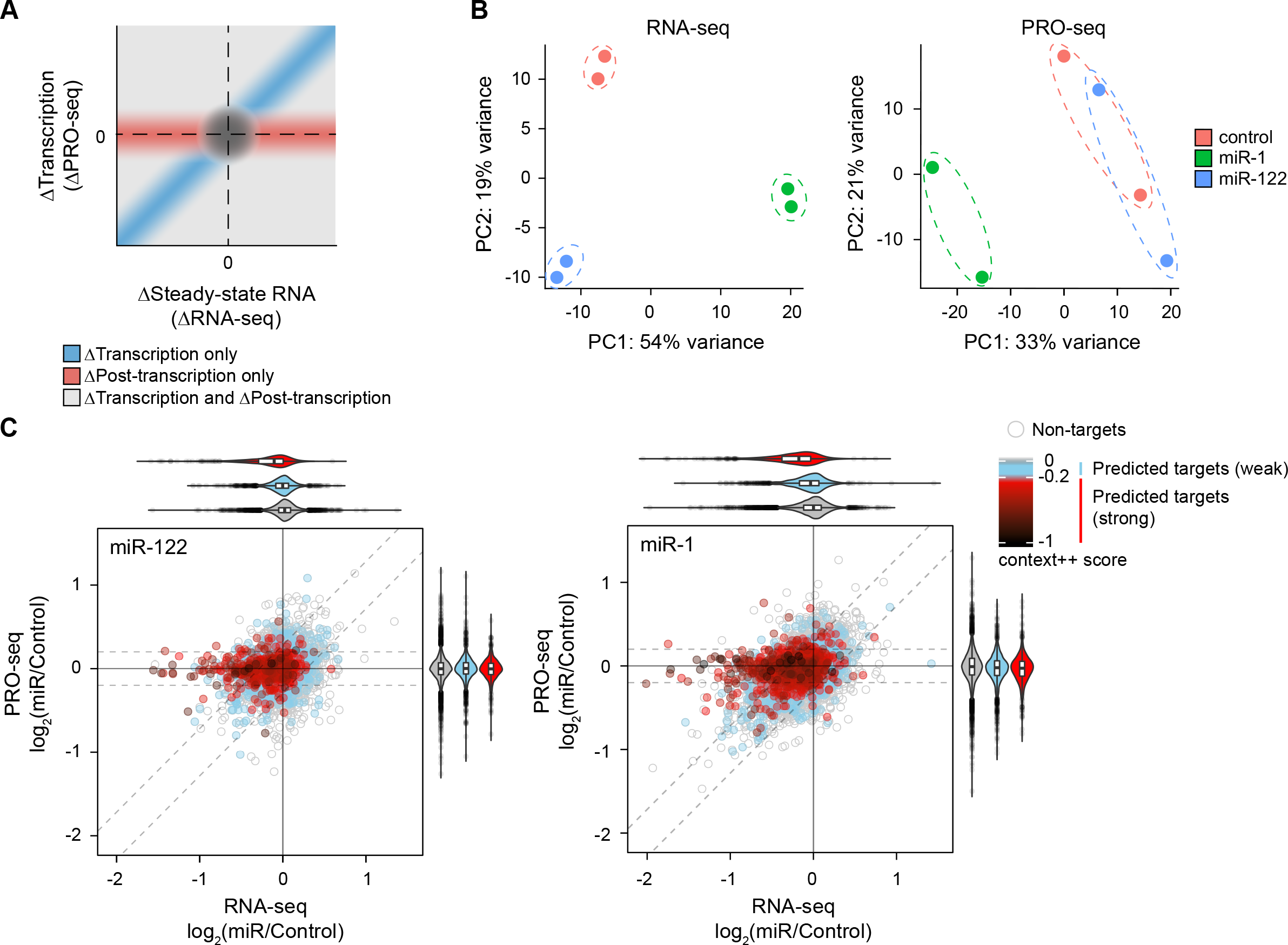
Combined analysis of RNA-seq and PRO-seq identifies genes subject to post-transcriptional regulation. (**A**) Schematic representation of combined analysis of RNA-seq and PRO-seq data. Changes in transcriptional output (y-axis; PRO-seq) and mRNA abundance (x-axis; RNA-seq) are plotted. (**B**) Principal component analysis of RNA-seq (left) and PRO-seq (right) data, for cells induced with miR-1 or miR-122, and control cell lines without induced miRNAs. (**C**) Dot plots depicting changes in steady-state mRNA levels (RNA-seq; x-axis) and transcriptional output (PRO-seq; y-axis) for all expressed genes (dots) upon miR-122 (left) or miR-1 (right) expression. The grey, blue, and red dots represent genes without predicted target sites, predicted weak targets and predicted strong targets of cognate miRNAs, respectively. The gradient of red color indicates predicted efficacy of targets (TargetScan context++ score). A log_2_ fold change of 0.2 in PRO-seq (Δtranscription) and CARP (Δpost-transcription) is indicated using horizontal and diagonal dotted lines, respectively. The violin plots on the top and right illustrate distributions of log_2_ fold-changes in RNA-seq and PRO-seq, respectively, for the different categories of genes, color-coded as described above.

To simultaneously measure transcriptional and post-transcriptional changes upon miRNA induction, we performed PRO-seq and RNA-seq on cells expressing miR-1 or miR-122 and compared them to control cells lacking a miRNA hairpin. Principal component analysis (PCA) of the RNA-seq data showed that replicate transcriptomes cluster tightly, indicating high data reproducibility, and that the transcriptomes of cells expressing miR-1 and miR-122 are distinct from each other and from the control cells (Figure 1B, left panel). When we performed PCA of the PRO-seq data, however, we found that the genome-wide transcriptional profile of control and miR-122 expressing cells were not well separated (Figure 1B, right panel), perhaps indicating that miR-122 does not elicit wide-spread changes in transcription. Indeed, transcription of 370 genes were significantly altered (q-value < 0.05) in response to miR-1, whereas only 9 genes were significantly altered in response to miR-122, supporting this interpretation (Supplementary Figure S1H).

To investigate the effects of miRNA induction on transcriptional and post-transcriptional regulation, we compared changes in PRO-seq and RNA-seq across individual genes. The rates of synthesis and degradation together determine steady-state mRNA levels. Therefore, we reasoned that changes in PRO-seq signal (ΔPRO-seq), representing changes in transcriptional outputs, subtracted from changes in RNA-seq (ΔRNA-seq), representing changes in steady-state RNA levels, would provide a quantitative readout for post-transcriptional regulation. This combined analysis of RNA-seq and PRO-seq, referred to as CARP hereafter, identified many genes subjected to post-transcriptional repression in response to miR-122 or miR-1, as indicated by repression in RNA-seq profiles without any changes in PRO-seq (Figure 1C). In fact, the 3UTRs of most of these genes contained target sites predicted to be strongly effective, as defined by TargetScan ((16); context++ score < −0.2; referred to as predicted strong targets hereafter). Correlating post-transcriptional repression with predictions of site efficacy, most genes containing target sites predicted to be weakly effective (as defined by TargetScan context++ score > −0.2; referred to as predicted weak targets hereafter), showed very subtle changes, if any, in RNA-seq compared to genes without predicted target sites (Figure 1C; log_2_ fold-change of - 0.02 and −0.03 for miR-122 and miR-1, respectively). Additionally, many genes, including predicted strong targets, demonstrated concordant changes in both PRO-seq and RNA-seq in cells expressing miR-1 (Figure 1C, right panel; Pearson correlation, r=0.42), likely representing genes that are regulated predominantly at the level of transcription, and not direct targets of miR-1. However, such concordant changes were minimal in cells expressing miR-122 (Figure 1C, left panel, r=0.19), consistent with the PCA of PRO-seq profiles (Figure 1B, right panel). Furthermore, the majority of genes were insensitive to induced miRNAs (63% and 79% genes in miR-1 and miR-122 samples, respectively, with absolute log_2_ fold-change smaller than 0.2 in both RNA-seq and PRO-seq), as expected for single miRNA perturbation experiments. These unchanged genes included 45% and 60% of predicted targets (including strong and weak) of miR-1 and miR-122, respectively (47% and 55%, if considering conserved target sites predicted by TargetScan, at a probability of conserved targeting threshold > 0.5), likely representing high false-positive rates of prediction algorithms, as reported previously (4,8,19,60). Taken together, CARP robustly deconvolutes and quantifies post-transcriptional and transcriptional regulation, enabling a robust experimental framework for distinguishing the direct miRNA targets from the resulting downstream regulatory changes.

### Combined analysis of RNA-seq and PRO-seq identifies direct miRNA targets

Direct miRNA targets correspond to transcripts bound by miRNA-loaded AGO at target sites, resulting in accelerated decay and/or translational repression. To assess the ability of CARP to detect direct targets, we first performed a likelihood ratio test (LRT) (43) to identify genes that experience a significant post-transcriptional change – that is, a significant change in steady-state mRNA level after accounting for any change in transcription. Consistent with the role of a miRNA as a negative regulator, most of the genes subject to post-transcriptional regulation (98% and 75% for miR-122 and miR-1, respectively) exhibited repression upon miRNA induction (Figure 2A and B). To evaluate further the properties of post-transcriptionally regulated genes, we assessed the presence of predicted target sites for the cognate induced miRNAs, using TargetScan v7.0 (16). The 3’UTRs of the majority of post-transcriptionally down-regulated genes contained predicted target sites for miR-122 or miR-1 (90% and 63%, respectively), although we note that many additional predicted targets (567 and 2,573 predicted strong and weak targets, respectively, for miR-122; and, 742 and 1,969, respectively, for miR-1) were not detectably repressed. To confirm that the post-transcriptional repression of the predicted targets (potential direct targets) is due to direct binding of miRNA-loaded AGO, we performed UV crosslinking and immunoprecipitation of AGO followed by sequencing (eCLIP-seq; (29)) to identify mRNAs bound by AGO. We reasoned that if the post-transcriptionally repressed genes containing predicted target sites are indeed direct targets, their 3UTRs should exhibit increased binding of AGO in cells expressing miR-122 or miR-1 compared to control cells. Following quality trimming and mapping of eCLIP data, peaks of eCLIP reads, representing regions of the transcriptome occupied by AGO, were identified using CLIPper (50). Although AGO binds predominantly at miRNA target sites in 3UTRs (21), a majority (53%) of eCLIP peaks overlapped with introns, with only 17% peaks in 3 UTR (Supplementary Figure S2A), implying high levels of background in the eCLIP dataset. Further examination of the peak counts revealed that the replicates exhibited large variations (Supplementary Figure S2B), a limitation associated even with more mature AGO CLIP protocols. To remove irreproducible peaks, we calculated irreproducibility discovery rate (IDR; (51)), a method appropriate for eCLIP data (29), and filtered peaks with less than 0.1 IDR, resulting in an average of 5,192 peaks per sample (Supplementary Figure S2C). A majority (54%) of our reproducible peaks overlapped with 3UTRs, with only 7% in introns (Supplementary Figure S2D), suggesting that the intronic peaks found in our unfiltered data and in previous reports (24) likely correspond to background signal. To assess the quality of our eCLIP data further, we compared relative AGO density (see methods) and post-transcriptional repression of predicted targets as a function of predicted target site efficacy (TargetScan) (16). As the predicted site efficacy increased, we observed an increase in relative AGO density in 3UTRs concomitant with an increase in post-transcriptional repression (Supplementary Figure S2E), indicating that our filtered eCLIP data reflect high quality AGO density profiles.

**Figure 2.**
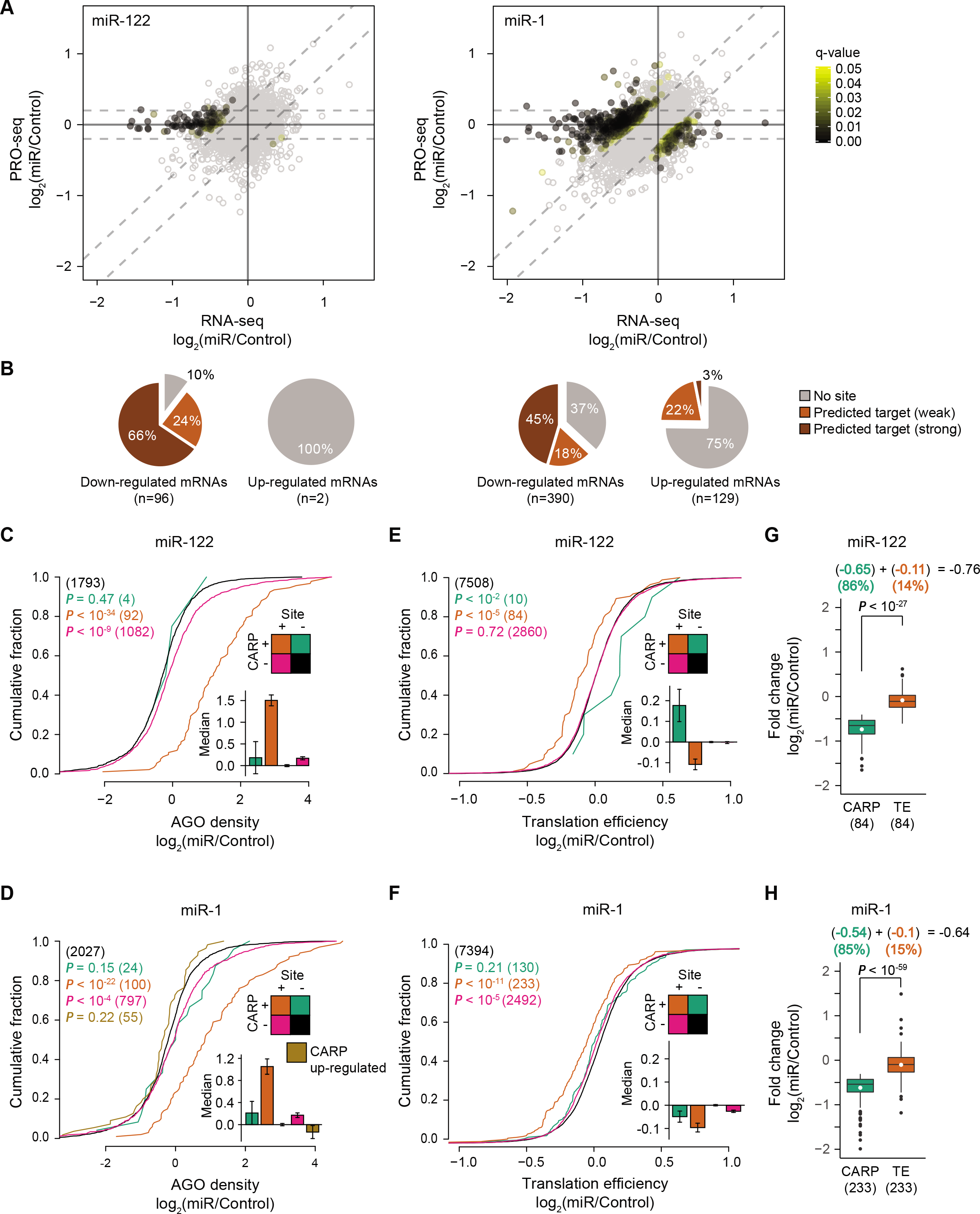
Identification and analysis of direct miRNA targets. (**A**) Identification of post-transcriptional changes in cells expressing miR-122 (left) or miR-1 (right). The dot plots are as described in Figure 1C, except that the filled and open dots illustrate genes with and without significant post-transcriptional change (q-value threshold of 0.05), respectively. Filled dots are color-coded based on q-value. (**B**) Pie charts depicting the percentages of genes without predicted target sites (grey), genes predicted to be weak targets, and genes predicted to be strong targets for post-transcriptionally down-regulated and up-regulated sets of genes in cells expressing miR-122 (left) or miR-1 (right). (**C** and **D**) Cumulative distribution plots of relative AGO density at eCLIP peaks within 3’UTRs in cells expressing miR-122 (C) or miR-1 (D) compared to the control cells. Genes were partitioned into four color-coded groups based on presence or absence of predicted target site (Site), and whether genes are significantly post-transcriptionally repressed in response to miRNA induction (CARP). Genes without predicted target sites and not subject to post-transcriptional regulation (Site-/CARP-; black) was used as the background set and compared to the remaining three groups. P values and number of eCLIP peaks (in parentheses) are indicated by color (see legend). A bar plot with standard error (error bars) is shown within each cumulative distribution plot depicting median log_2_(miR/Control) AGO density (y-axis) for each category. (**E** and **F**) Cumulative distribution plots of translational efficiency in miR-122 (E) or miR-1 (F); otherwise as described in panels C and D. P values and number of genes (in parentheses) are indicated by color. A bar plot is shown within each cumulative distribution plot depicting median log_2_(miR/Control) translational efficiency (y-axis) for each category. (**G** and **H**) Box plots comparing contributions of mRNA degradation (green) and translational inhibition (orange) in miRNA-mediated post-transcriptional repression of CARP-identified direct targets in cells expressing miR-122 (G) or miR-1 (H). The plotted log_2_ fold-changes were normalized by the median regulation observed for the background genes (genes lacking predicted 3UTR sites for the cognate miRNA that showed no significant change in respective measurements). Mean and median log_2_ fold changes are indicated by white point and horizontal bar, respectively. The medians of log_2_ fold changes were used to quantify relative contributions of decay and translational repression (see equation above each plot).

Using the filtered high quality AGO eCLIP data, we asked whether our classification of predicted targets exhibiting significant post-transcriptional repression as direct targets is supported by AGO binding. We found that relative AGO density is strongly enhanced in 3UTRs of post-transcriptionally down-regulated genes containing predicted target sites, with 85 and 90% of genes bound by more AGO in cells expressing miR-122 and miR-1, respectively, compared to control cells (Figure 2C and D, orange line). Furthermore, consistent with the absence of significant post-transcriptional repression for a large number of predicted targets, we observed a negligible, albeit statistically significant, increase in AGO density in their 3’UTRs (Figure 2C and D, pink line). Collectively, these observations indicate that the intersection of genes distinguished as post-transcriptionally repressed by CARP with those also containing predicted target sites represents a set of high confidence set of direct miRNA targets, and that CARP robustly distinguishes these direct targets from the large numbers of predicted yet ineffective targets.

The robustness of CARP is dependent on its ability to reliably profile post-transcriptional regulation. To evaluate the specificity of post-transcriptional changes identified using CARP, we assessed the overlap between miR-1-specific and miR-122-specific post-transcriptional changes. Since the seed sequences of these miRNAs differ by only a single nucleotide (miR-122: GGAGUGU; miR-1: GGAAUGU), they can act as negative controls for each other. We found that only five genes were in common between the genes repressed post-transcriptionally by miR-1 and miR-122, all of which contained predicted target sites for both miRNAs in their 3UTRs (Supplementary Figure S2F). There was no overlap for the post-transcriptionally up-regulated genes. To further evaluate the reliability of CARP, we sought to investigate whether the targets we identified in HEK293 cells correspond to targets regulated by these miRNAs *in vivo*. We reasoned that the true targets of tissue-specific miRNAs such as miR-122 (liver-specific) and miR-1 (muscle-specific) would be expressed at lower levels in those specific tissues, a defining feature of tissue-specific miRNAs (52,61–63). To this end, we compared the expression of miR-1-specific and miR-122-specific direct targets identified by CARP across 53 different tissues from Genotype-Tissue Expression (GTEx) Project (BROAD Institute). Consistent with our hypothesis, only the liver and skeletal muscle tissues exhibited significantly lower expression of CARP-identified direct targets of miR-122 and miR-1, respectively, compared to the randomly selected set of genes (Supplementary Figure S2G). Collectively, these results establish that CARP detects true targets of miRNAs with strong specificity.

CARP relies on mRNA steady-state measurements, and therefore cannot detect miRNA targeting that results exclusively in translational repression. Thus, we investigated whether predicted targets that lacked significant post-transcriptional regulation (detected by CARP) were undergoing translational repression. To address this possibility, we performed ribosome profiling (30) to measure changes in translation upon miRNA induction, comparing cells expressing miR-1 or miR-122 to control cells. We first evaluated the change in translational efficiency (see methods) for all genes in miRNA-expressing cells compared to control cells. We found that only a small number of predicted targets (including strong and weak) exhibited significant evidence of translational repression (Supplementary Figure S2H; 8 and 25 genes for miR-122 and miR-1, respectively). Of these translationally repressed targets, a subset also exhibited accelerated mRNA decay (2 and 10, for miR-122 and miR-1, respectively), whereas others (6 and 15) were repressed exclusively via translational regulation (Supplementary Figure S2H). We next assessed the translational efficiency of CARP-identified direct targets, which revealed that these miRNA targets experienced significant, albeit subtle, translational repression compared to non-targets (Figure 2E and F, compare orange lines to black lines). These results indicated that miRNAs expressed at physiological levels repress targets predominantly via accelerated mRNA decay and that targets regulated principally by translational regulation are exceedingly rare; thus, CARP is well suited for identifying direct targets.

Excluding certain specific cellular contexts (64,65), mRNA decay is thought to be the major contributor of miRNA-mediated repression, with translational repression playing only a minor role (66). To revisit this important question with miRNAs expressed at physiological levels, we next used our ribosome profiling dataset to quantify contributions from both translational repression and mRNA decay in miRNA-mediated repression. We found that the magnitude of mRNA decay was significantly greater than that of translational repression (6-fold greater for both miR-122 and miR-1; Figure 2G and H). We found that reduced mRNA levels explain most of the overall regulation observed (86% and 85%, for miR-122 and miR-1, respectively) compared to translational repression (14% and 15%, Figure 2G and H). These estimates are consistent with those obtained in previous studies using transiently transfected miRNAs (67), and in marked contrast with other reports (68–70).

### Combined analysis of RNA-seq and PRO-seq facilitates discovery of indirect targets

In addition to direct targets, we also found a smaller portion (10% and 37%, for miR-122 and miR-1, respectively) of post-transcriptionally repressed genes that lacked predicted target sites (Figure 2 B). This set represents either direct targets containing non-canonical target sites, or indirect targets of miRNAs for which the regulation is itself post-transcriptional. Notably, the absence of significantly increased relative AGO density within the 3’UTRs of these genes (Figure 2C and D; compare green lines to black) indicates that they are indirect targets of miR-122 and miR-1, which are down-regulated post-transcriptionally. Additionally, we observed post-transcriptional up-regulation of 129 genes upon miR-1 induction, which likely represent post-transcriptional indirect targets of miR-1, as miRNAs are repressive molecules (Figure 2B). Consistent with this interpretation, most (75%) up-regulated genes lacked predicted target sites, and the up-regulated gene set also lacked an increase in relative AGO density (Figure 2D). Thus, while we initially assumed that the indirect targets must be primarily regulated at the level of transcription, our results indicate that indirect targeting triggered by miR-1 results in both widespread transcriptional and post-transcriptional regulation. Presumably, this broader indirect regulation is a result of miRNA-mediated repression of mRNA transcripts coding for transcription factors and regulatory RNA-binding proteins, respectively. Indeed, the direct targets of miR-1 included many genes coding for transcription factors (55) and RNA binding proteins (71), whereas miR-122 repressed relatively few (Supplementary Figure S2I). Taken together, these data indicate that while the regulatory network of miR-122 is comprised primarily of direct targets in HEK293 cells, miR-1 elicits more complex responses involving both direct and indirect targets to regulate gene expression in these cells. In addition, a substantial proportion of the indirect effects induced by miR-1 occurs via post-transcriptional regulation. Importantly, however, the lack of predicted target sites within such genes enables reliable partitioning of post-transcriptionally repressed genes into those that are direct targets and those that are indirectly regulated by additional miRNA-independent decay pathways. In addition, we note that, because our approach directly quantifies transcription and mRNA levels, CARP is well-suited to disentangling these complex transcriptional and post-transcriptional regulatory changes.

### Partitioning modes of regulation elicited by miRNAs

To systematically investigate the utility of CARP, we examined relationships among three categories of genes: (i) genes exhibiting reduced mRNA abundance in response to either miR-122 or miR-1, which we measured using RNA-seq, (ii) genes predicted to be strong microRNA targets, which we determined using TargetScan (context++ < −0.2) (16), and (iii) genes that we found to undergo significant post-transcriptional repression in response to miR-122 or miR-1. We compared these three gene sets, and depicted the results using Venn Diagrams (Figure 3A). A substantial fraction of predicted targets, for both miR-122 and miR-1 (72 and 57%, respectively), do not exhibit evidence of significant changes in mRNA abundance or in post-transcriptional levels, consistent with established high rates of false positive predictions (set *a* in Figure 3A). Consistent with this interpretation, this set of genes exhibited no change in CARP, RNA-seq, PRO-seq or ribosome profiling (Figure 3B). Similarly, we found minimal evidence for increased AGO density in the 3UTRs of this gene set (Figure 3 C and D, Supplementary Figure S3A). We note that the mRNA transcripts of the majority (54 and 57% for miR-122 and miR-1, respectively) of these genes exist as 3UTR isoforms that contain the predicted target sites; thus, the absence of regulation cannot be attributed solely to alternative processing (Supplementary Figure S3B). Taken together, these results indicate that this category likely corresponds to false-positive predictions and/or targets that are not effective in this cell line.

**Figure 3.**
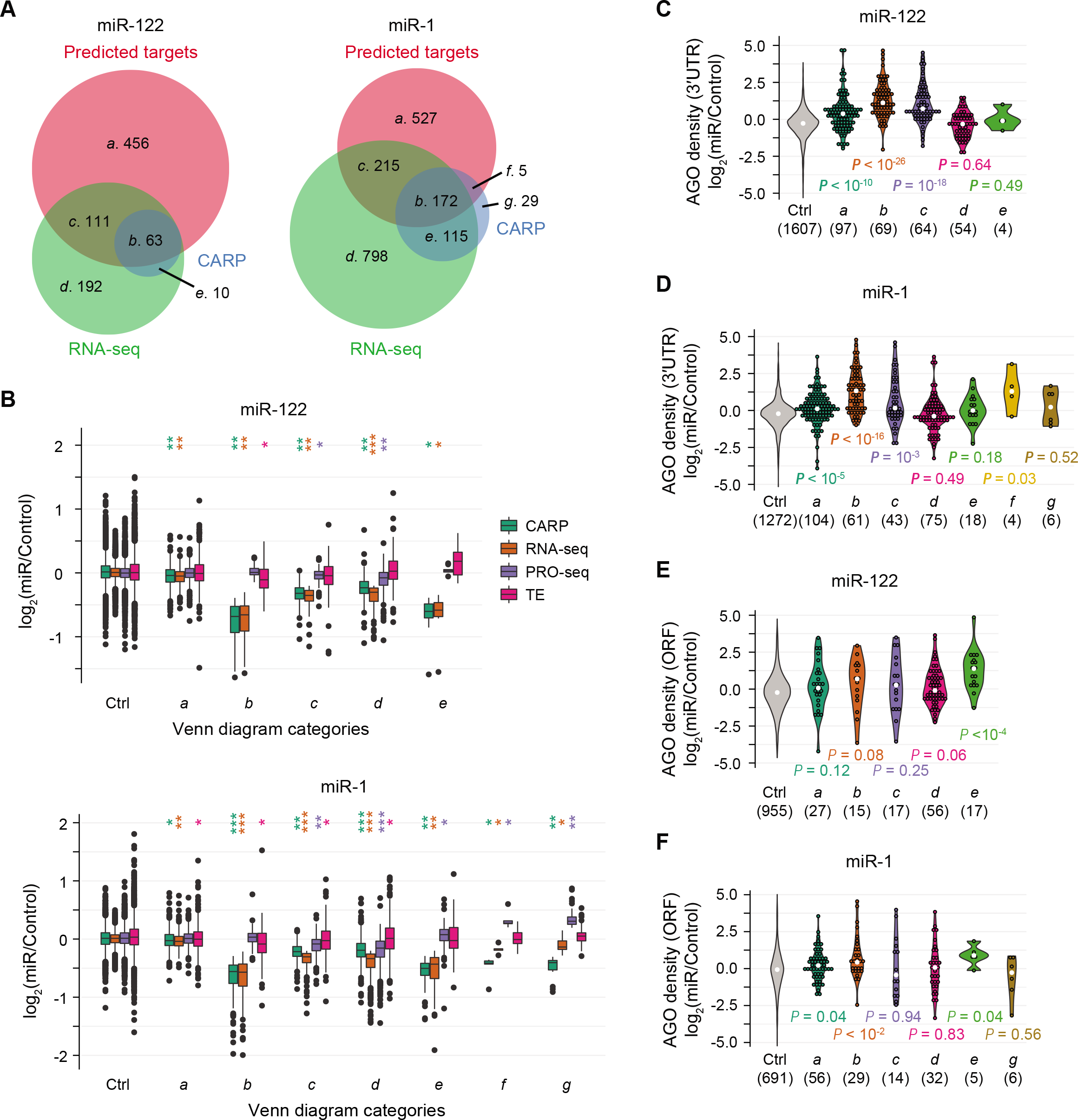
CARP enables discovery of different miRNA regulatory mechanisms. (**A**) Venn diagram analysis of predicted strong targets of cognate miRNAs (TargetScan; context++ score < −0.2; red set), genes experiencing reduced transcript levels (RNA-seq; q-value < 0.05 and log_2_(miR/Control) < −0.2; green set) and post-transcriptional repression (CARP; q-value < 0.05 and log_2_(miR/Control) < −0.2; blue set) in response to miR-122 (left) or miR-1 (right). Genes expressed at low levels (reads per million ≤ 1 in any sample of RNA-seq or PRO-seq datasets) and the predicted weak targets (context++ score > −0.2) were excluded from this analysis. Geneset labels (*a*-*g*) and number of genes in each set are displayed. Areas are proportional to numbers of genes in each set. (**B**) Boxplots illustrating distributions of log_2_(miR/Control) for indicated gene-sets for post-transcriptional regulation (CARP; green), mRNA abundance (RNA-seq; orange), transcriptional output (PRO-seq; purple) and translational efficiency (TE; pink) in cells expressing miR-122 (top) or miR-1 (bottom). The control set (Ctrl) consists of genes that lack significant change in RNA-seq and CARP (q-value > 0.05) whose 3UTRs are devoid of predicted target sites. Gene-sets *a* through *g* correspond to gene-sets depicted in panel A. P values comparing each gene-set to the Ctrl are indicated as follows: *P < 0.01, **P < 10^-10^, ***P < 10^-100^. (**C** and **D**) Violin plots depicting distributions of relative AGO density at eCLIP peaks within 3UTRs of gene-sets *a*-*g* (as in panel A) in cells expressing miR-122 (C) or miR-1 (D). The control (Ctrl) gene-set is as described in panel B. The numbers in parentheses (x-axis) indicate numbers of eCLIP peaks found within 3’UTRs of each gene-set. White dots represent median values and colored dots represent individual values of the distributions. P values comparing each set to the Ctrl are indicated. (**E** and **F**) Violin plots identical to those in panels C and D except depicting AGO density at eCLIP peaks within ORFs.

The second subset of genes we considered were those that contain predicted target sites whose mRNA abundance is reduced in response to the cognate miRNA, and which we determined were regulated post-transcriptionally (set *b*, Figure 3A). Importantly, this gene set exhibited the largest reduction in mRNA levels in response to the cognate miRNA (Figure 3B). Moreover, post-transcriptional regulation was the primary (and for many genes, sole) factor responsible for repression (Figure 3B). Congruently, this set of genes also demonstrate the strongest increase in relative AGO density in their 3UTRs in response to the induced miRNAs (Figure 3C and D, Supplementary Figure S3A). This target set corresponds to direct miRNA targets. We note that translational repression for these set of targets is minimal (Figure 3B).

We next considered predicted target genes that were down-regulated in mRNA abundance but without significant post-transcriptional repression (set *c*, Figure 3A). As a set, such genes exhibit minimal evidence of either miRNA-induced post-transcriptional repression or translational repression (Figure 3B). We recognize that a subset of these genes may represent *bona fide* direct targets, but with reduced site efficacy, thus reducing the ability to detect experimentally.

Nevertheless, the average post-transcriptional repression is substantially less (Figure 3B) than that observed for direct targets defined by CARP (set *b*). Moreover, although relative AGO density is significantly increased for these genes, the average magnitude of this increase is minimal (1.6 and 1.1-fold, compare to 2.2 and 2.5-fold for set *b*; for miR-122 and miR-1, respectively; Figure 3C and D, Supplementary Figure S3A). It is important to note that in numerous conventional analyses of miRNA target sets, including our previous work (72), all genes in set *c* would have been erroneously declared as direct targets.

An important class of genes are those that have lower steady-state mRNA levels but do not contain predicted target sites for miR-122 or miR-1 (set *d*, Figure 3A). Similar to set *c*, they exhibit a negligible magnitude of post-transcriptional regulation (1.17 and 1.14-fold for miR-122 and miR-1, respectively) even though mRNA levels are significantly reduced in response to the induced miRNAs. Concordantly, in response to miR-1 or miR-122, we did not observe any significant increase in relative AGO density in 3’UTRs or in ORFs of these genes (Figure 3C-F, Supplementary Figure S3A and C), nor any evidence of translational repression (Figure 3B), ruling out the possibility that they are potential direct targets containing non-canonical target sites. Interestingly, these genes also demonstrated subtle repression in transcriptional output (Figure 3B, PRO-seq). Perhaps this set represents targets of feed-forward control, which are repressed minimally at both a transcriptional and post-transcriptional level, to ultimately exhibit effective repression in mRNA abundance. As a group, however, they exhibit characteristics indicative of indirect miRNA targets regulated minimally at both transcriptional and post-transcriptional levels.

We also identified genes that are significantly post-transcriptionally repressed, but which lack predicted miRNA target sites (set *e*, Figure 3A). The magnitude of post-transcriptional repression for this gene set matched set *b*, the set of confidently identified direct targets of miR-122 and miR-1. We considered two possibilities to explain the observed post-transcriptional repression for set *e*: either they represented indirect targets whose regulation is post-transcriptional, or they represent direct targets lacking conventional miRNA target sites. We recognize that both scenarios might apply to different genes within the set. Notably, we did not observe increased relative AGO density in the 3’UTRs of these genes (Figure 3C and D), consistent with lack of predicted 3’UTR target sites. Given the absence of increased relative AGO density in the 3’UTR, we considered whether these genes might be repressed via target sites within their ORFs. For miR-1, we detected only five AGO eCLIP peaks in ORFs of 115 genes of set *e* and the increase in relative AGO density at these peaks was very weak (Figure 3F), implying that these genes are indeed indirect targets of miR-1, which are regulated post-transcriptionally, and independent of the miRNA pathway. We note that this interpretation is consistent with our observation that many genes were post-transcriptionally up-regulated in response to miR-1; that is, miR-1 induces widespread indirect post-transcriptional regulation. In contrast, we observed a strong increase in relative AGO density in the ORF of miR-122-regulated genes in set *e* (Figure 3E, Supplementary Figure S3C), comparable in magnitude to that observed in the 3UTR of *bona fide* direct targets (set *b*; 2.6 and 2.2-fold increase in ORFs of set *e* genes and in 3UTRs of set *b* genes, respectively). We note that set *e* encompasses only few (n=10) genes in response to miR-122. Nevertheless, these results suggest that miR-122 directly regulates a small group of genes via target sites in the ORF.

In response to miR-1, but not miR-122, we observed a small number of genes that are both predicted strong targets and significantly repressed post-transcriptionally, but without a detectable change in mRNA abundance (Figure 3A and B; set *f*). We hypothesized that these genes are direct targets, but also transcriptionally up-regulated, resulting in no net change in mRNA levels. Consistent with this idea, we observed a strong increase in relative AGO density, comparable to that seen in set *b*, in 3UTRs for such genes (Figure 3D; average fold changes of 2.51 and 2.45, for sets *b* and *f*, respectively). The weaker statistical significance of relative AGO density is due to the small number of genes in this category. Importantly, we also observed increased transcription of these genes (Figure 3B), confirming transcriptional up-regulation triggered by miR-1. This gene set illustrates an interesting class of direct targets that are invisible to studies reliant on RNA-seq alone.

The final possible class of genes corresponds to those exhibiting post-transcriptional repression without any significant change in mRNA abundance and without predicted target sites (Figure 3A and B, set *g*). We observed this class of genes in response to miR-1 alone. Consistent with the absence of predicted target sites, we did not observe an increase in relative AGO density in their 3UTRs (Figure 3D, Supplementary Figure S3A), nor within ORFs of these genes (Figure 3F, Supplementary Figure S3C), suggesting that these genes are indirect targets that are repressed post-transcriptionally reminiscent of the genes in set *e*. Similar to set *f*, these genes also exhibited transcriptional up-regulation, the impact of which is masked by post-transcriptional repression, resulting in no change in mRNA abundance (Figure 3B). Collectively, this set of genes represents indirect targets that are regulated at both the transcriptional and post-transcriptional levels.

It is important to acknowledge that partitioning genes into sets and subsets according to RNA-seq and PRO-seq signals necessitates use of statistical thresholds. Accordingly, we examined whether our interpretations (relating to Figure 3A) are robust over a range of reasonable thresholds. Overall, our observations remain consistent across a series of statistical thresholds (Supplementary Figure S3A and C). For example, irrespective of the significance threshold, we observed large numbers of predicted strong targets which did not exhibit evidence of post-transcriptional repression, consistent with high rates of false-positive predictions. Notably, consistent with CARP’s ability to robustly separate true direct targets (set *b*) from false targets of conventional approaches (set *c*), even when we assessed the most lenient threshold (q-value < 0.1), we found many predicted targets that exhibited reduced mRNA abundance without a significant change in post-transcriptional regulation (set *c*). Taken together, our approach identifies not only the robust direct targets but also discovers a variety of complex regulatory outcomes elicited by miRNAs.

### MicroRNA-specific targeting of sites located in open reading frames

The prevalence and efficacy of miRNA target sites within coding sequence is unclear and controversial. Whereas certain studies indicate that such sites exert a negligible effect, perhaps due to ribosome□mediated removal of miRNA-loaded AGO from translating ORFs (73), others indicate that such sites are often effective in mediating repression (74). Here, we revisited this important question, using our improved ability to quantify post-transcriptional regulation.

We observed increased relative AGO density within ORFs in response to miR-122 for a small set of genes (Figure 3E, Supplementary Figure S3C; set *e*); importantly, this set of genes was devoid of predicted 3’UTR sites, and exhibited no evidence of increased relative AGO density within the 3’UTR (Figure 3C, Supplementary Figure S3A; set *e*). In addition, this set of genes was repressed post-transcriptionally in response to miR-122; thus, it seemed plausible that miR-122 was directly repressing mRNAs of these genes using target sites within the ORFs. To systematically examine miRNA-mediated regulation of ORF target sites, we grouped genes based on the location of potential target sites (matches to 8mer, 7mer-m8 and 7mer-A1 site motifs) within the mRNA and compared the degree of post-transcriptional regulation. In response to miR-122, genes containing predicted miR-122 target sites in the ORF were significantly repressed, and the average efficacy of these sites exceeded the efficacy of those predicted to be weakly effective when located within 3UTRs (Figure 4A, compare orange line with purple; P-value < 10^-6^). For miR-122, we also found that post-transcriptional repression grows stronger with increasing number of miR-122 target sites in ORFs (Supplementary Figure S4A). Consistent with activity of miR-122 ORF sites, we observed increased relative AGO density in ORFs containing miR-122 target sites (Supplementary Figure S4B). Moreover, we observed that sequence conservation of predicted miR-122 ORF sites is associated with the site efficacy, with conserved ORF sites exhibiting 2.7-fold stronger repression compared to all ORF sites, suggesting biological importance of such sites (Supplementary Figure S4C). We note that the efficacy of predicted conserved ORF sites is weaker compared to predicted conserved 3UTR sites (Supplementary Figure S4C). In stark contrast, we found no evidence for effective miR-1 target sites within ORFs (Figure 4A), even when we examined genes harboring multiple miR-1 ORF sites or conserved miR-1 ORF sites (Supplementary Figure S4A and C, respectively), suggesting that specific miRNAs differ in their ORF targeting propensities. Sites within 5UTRs were ineffective for both miRNAs (Figure 4A).

**Figure 4.**
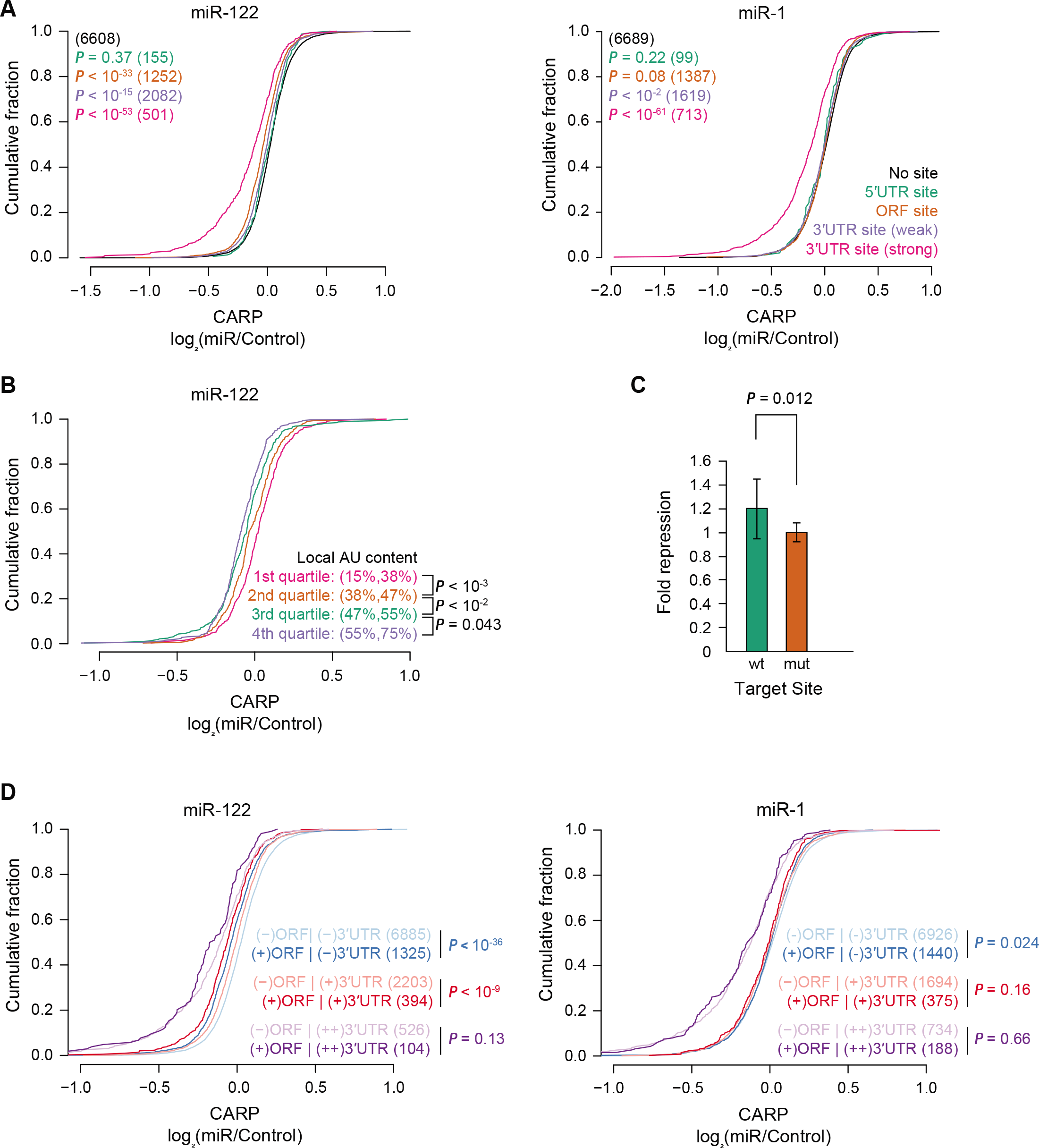
MicroRNA-specific targeting of sites located in open reading frames. (**A**) Cumulative distribution plots of post-transcriptional regulation (CARP, x-axis) of genes in response to miR-122 (left) or miR-1 (right). Distributions for genes containing predicted sites (matches to 8mer, 7mer-m8 or 7mer-A1 site motifs) located in the 3’UTR, ORF or 5’UTR, or with no site are plotted. Genes with sites in multiple regions are excluded from this analysis. The genes containing predicted 3 UTR sites were divided into two groups according to predicted efficacy of targets (predicted weak targets, context++ > −0.2; predicted strong targets, context++ < −0.2). Each group of genes containing target sites were compared to the control group without any predicted sites. P values and number of genes (in parentheses) for each category are indicated by color. (**B**) Cumulative distribution plot comparing the post-transcriptional regulation of miR-122 targets partitioned by local AU content (fraction of As and Us in 30 nucleotides) around the predicted ORF sites. The range (in percentage) of local AU content for each quartile are presented in parentheses. The observed regulation for each pair of consecutive quartiles was compared. (**C**) Luciferase reporter assay showing miRNA-mediated repression of a reporter containing a candidate miR-122 ORF target site with 78 nucleotides of endogenous coding sequence cloned in-frame at the C-terminus of firefly luciferase. After normalizing to the transfection control, each wild-type (wt) and mutant (mut) target site reporter co-transfected with a miR-122 mimic was normalized to those co-transfected with a non-specific miRNA mimic. These normalized values are plotted, with error bars representing standard deviation (n=12). (**D**) Cumulative distribution plots of post-transcriptional regulation (CARP; x-axis) for genes without predicted 3 UTR sites [(-)3UTR], predicted weak targets [(+)3UTR] and predicted strong targets [(++)3UTR], with [(+)ORF] or without [(-)ORF] predicted ORF sites. P-values were calculated for the indicated comparisons. The number of genes in each category are listed in parentheses.

Next, we investigated whether the miR-122 ORF sites share properties associated with functional 3UTR sites. Canonical 3UTR sites differ in their average efficacy depending on nucleotide sequence and pairing potential with the 5’ terminus of the miRNA (referred to as the miRNA seed (19)); we examined 8mer, 7mer-m8 and 7mer-A1 sites (ordered from stronger to weaker sites; (9)) located within ORFs. Additionally, we also inspected a set of non-canonical sites, referred to as G-bulged sites, that are subject to a marginal repression by miR-122(75). We found a consistent pattern for predicted miR-122 ORF sites, with highest average repression elicited by 8mer sites followed by 7mer-m8 and 7mer-A1 sites, with the least repression for G-bulged sites (Supplementary Figure S4D). Beyond the type of seed match, a major additional determinant of 3UTR target site efficacy is local AU content, with higher AU content correlating with increased strength of miRNA-mediated repression (9). We found that predicted miR-122 ORF sites embedded in AU-rich regions exhibit increased post-transcriptional repression, with average repression correlating with local AU content (Figure 4B). No such relationships for site type nor AU content were observed for miR-1 (Supplementary Figure S4D and E).

To validate the efficacy of predicted miR-122 ORF sites, we generated luciferase reporters containing translational fusions between luciferase and a short region (encoding 26 amino acids) of endogenous sequence containing a potential miR-122 site. We also generated negative control variants of these reporters, which contained two to four synonymous mutations within the target site, designed to inactivate the site. Reporter assays using the wild-type and mutant (inactive site) constructs in the presence of miR-122 or a control miRNA indicated repression of wild-type reporters in response to miR-122 (Figure 4C). However, only two of the ten sites tested mediated statistically significant repression (Figure 4C, Supplementary Figure S4F), consistent with the reduced efficacy of predicted ORF sites compared to predicted 3’UTR sites observed in genomewide analyses (Figure 4A and Supplementary Figure S4C). In agreement with the importance of AU content to miRNA-mediated repression, the observed repression in reporter assays correlated with local AU content, albeit not significantly, likely due to the limited sample size (Supplementary Figure S4G). These results provide additional evidence that miR-122 regulates post-transcriptional expression by targeting ORF sites.

Given the modest impact of ORF sites in post-transcriptional regulation, we wondered whether such sites might function in concert with 3’UTR sites. To answer this question, we evaluated three groups of genes, (i) genes without predicted 3 UTR sites, (ii) genes with predicted weak 3’UTR sites and (iii) genes with predicted strong 3’UTR sites, and asked if post-transcriptional regulation of these three groups changed depending on presence or absence of predicted ORF sites. Consistent with the efficacy of miR-122 ORF sites, for genes without miR-122 3’UTR sites, those that contained ORF sites were significantly down-regulated compared to those without (Figure 4D, left). Interestingly, genes containing predicted weak 3 UTR sites were significantly more repressed when their ORFs also contained miR-122 sites. In contrast, the presence of ORF sites in genes containing predicted strong 3 UTR sites did not provide additional benefit, particularly for those genes that already are strongly repressed (genes with log_2_(miR/Control) < −0.3). Consistent with our earlier results, we did not see any impact of miR-1 ORF sites on post-transcriptional regulation for genes with or without sites within the 3UTR (Figure 4D, right). Collectively, these observations suggest that miR-122 ORF sites potentiate miRNA-mediated repression for genes containing marginal sites within their 3UTR.

It has been suggested that effective miRNA target sites in ORFs elicit translational repression as a larger component of total repression, when compared to sites within 3UTRs (74). Our data, however, indicate that targeting of ORF sites by miR-122 mediated no significant effect on translation (Supplementary Figure S4H). To examine further whether translation status influences the degree of miRNA-mediated repression of ORF sites, we grouped genes based on the degree of translation, approximated using the ratio of ribosome profiling signal and expression in RNA-seq. We observed, irrespective of the amount of translation, a gradual increase in post-transcriptional repression with increasing number of predicted miR-122 ORF sites (Supplementary Figure S4I, left panel). Taken together, these results suggest that regulation triggered by miRNAs is mechanistically equivalent whether sites are located in the ORF or the 3UTR, and not detectably altered by the translation status of the transcript.

### Utility of CARP

To establish the general efficacy of CARP, we extended our approach to additional miRNAs. We reanalyzed our miRNA profiling data and selected five more miRNAs (miR-133a, miR-155, miR-302a, miR-372 and miR-373) whose expression is absent or negligible in HEK293 cells. We generated HEK293 cell lines stably expressing each of these miRNAs. Following induction of miRNAs for one week, we performed RNA-seq and PRO-seq to quantify the global changes in mRNA abundance and transcriptional output upon miRNA induction. Similar to the observations for miR-1 and miR-122, the predicted strong targets of these miRNAs exhibited reduced mRNA abundance without any change in transcription, and predicted weak targets experienced only a subtle change (Supplementary Figure S5A). Using the likelihood ratio test (43), we identified gene sets demonstrating significant changes in post-transcriptional levels and compared these genes with sets of predicted targets. The majority of post-transcriptionally down-regulated genes (63-97%) contained predicted target sites for the cognate miRNA, whereas a majority of the up-regulated genes lacked predicted target sites (Figure 5A), consistent with the suitability of CARP in estimating post-transcriptional regulation. Notably, the PRO-seq data revealed different degrees of altered transcription, with certain miRNAs eliciting widespread changes in transcription (Supplementary Figure S5B), which can be attributed to variable degrees of indirect targeting triggered by different miRNAs. These results indicate that CARP effectively deconvolutes post-transcriptional from transcriptional regulation.

**Figure 5.**
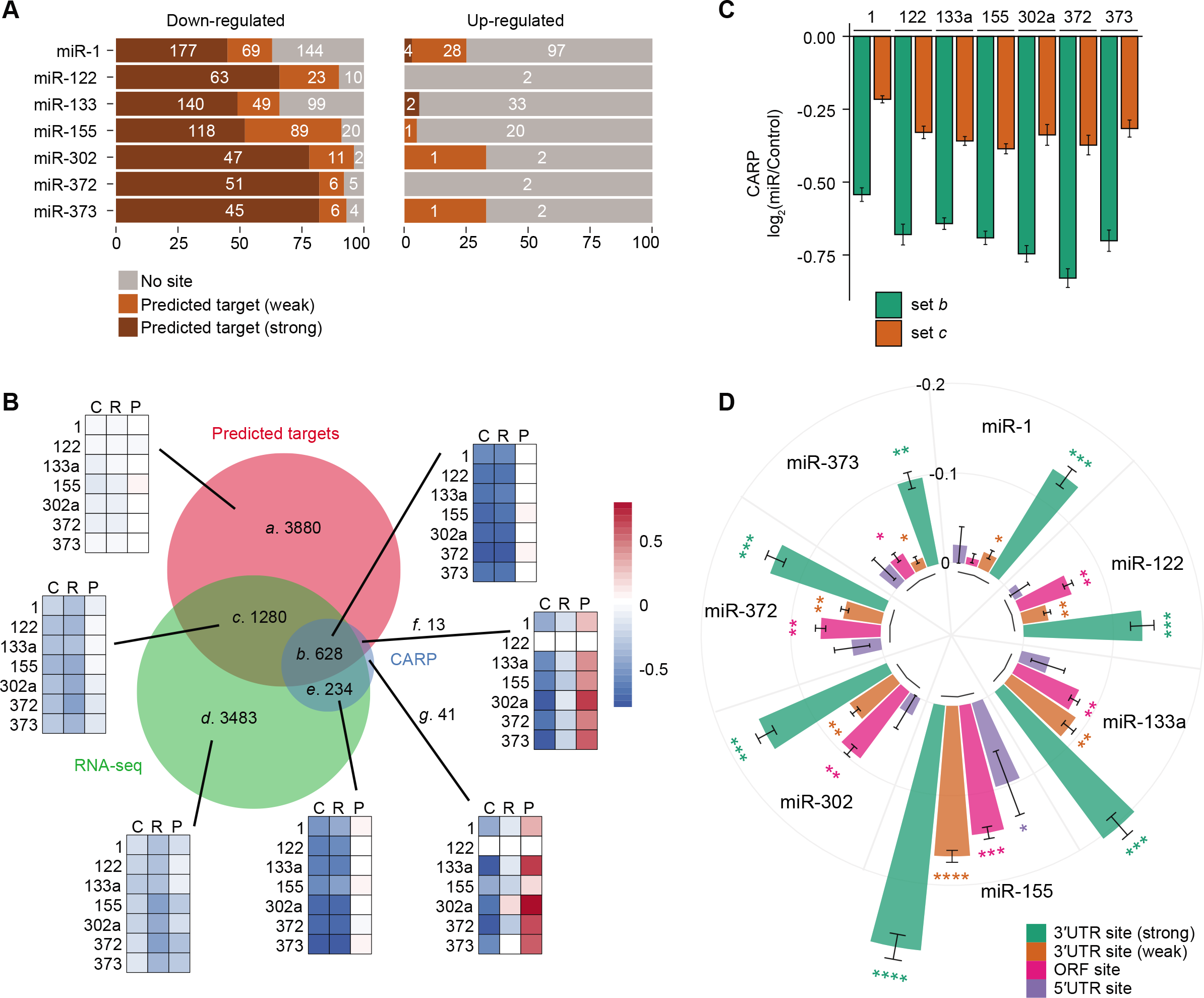
Utility of CARP. (**A**) Stacked bar plots of the fractions (x-axis) of genes without predicted target sites (grey), predicted weak targets (orange) and predicted strong targets (brown) in each of the two sets: post-transcriptionally down-regulated (left) and up-regulated (right) genes in response to different miRNAs (y-axis). The numbers of genes in each group are indicted. (**B**) Venn diagram with areas proportional to the number of genes in each set aggregated across all seven miRNAs, otherwise as described in Figure 3A. Heatmap depicts the median of log_2_(miR/Control) in post-transcriptional regulation (columns C, CARP), mRNA abundance (R, RNA-seq) and transcriptional output (P, PRO-seq) for each miRNA tested (rows). Scale bar denotes color code for log_2_(miR/Control). Heatmaps are shown for each gene-set of the Venn diagram. (**C**) Bar chart plotting post-transcriptional regulation (CARP, y-axis) for contextscore matched genes in set *b* (CARP+) and set *c* (CARP-). (**D**) Circular bar plot depicting median of log_2_(miR/Control) in post-transcriptional regulation in response to indicated miRNAs for genes containing predicted target sites as described in panel 4A, in either the 3UTR, ORF or 5 UTR. Genes containing 3UTR sites were divided into two groups according to predicted target site efficacy (strong, context++ score < −0.2; weak, context++ score > −0.2). The plotted median values were normalized by the median log_2_(miR/Control) of genes without predicted target sites (background set) to facilitate comparison between different miRNAs. Error bars represent standard error. P values comparing each plotted set with the corresponding background set are indicated as follows: *P < 0.01, **P < 10^-10^, ***P < 10^-50^, ****P < 10^-100^.

Next, we investigated relationships between the three categories of genes, as defined in Figure 3, for all 7 miRNAs examined in this study; thus, we partitioned genes by whether they were repressed post-transcriptionally, exhibited reduced mRNA levels, and contained a predicted strong miRNA target site. Consistent with results we observed for miR-1 and miR-122, a large number of predicted targets (set *a*) were not under the control of miRNAs (Figure 5B). Similarly, attributes of other sets of genes (sets *b*-*g*) were consistent with the results observed in Figure 3. In particular, direct miRNA targets identified by CARP (set *b*) exhibit strong post-transcriptional repression without any changes in transcription (Figure 5B, heatmaps for set *b*). The union of sets *b* and *c* represents hundreds of predicted targets that exhibit reduced mRNA abundance in response to the cognate miRNA, all of which would be considered as direct targets by conventional approaches. However, the improved resolution provided by CARP indicates that only a fraction (33%; set *b*) of those genes exhibit evidence of significant post-transcriptional repression, as opposed to only subtle changes for set *c* genes, for which transcriptional regulation (indirect regulation) is contributing to the observed reduction in transcript abundance (Figure 5B, heatmaps for set *c*). Consistent with this interpretation, the set *c* genes were predicted to contain target sites of somewhat lower efficacy than those in set *b* (Supplementary Figure S5C). Nevertheless, the set *b* genes exhibited far stronger post-transcriptional repression compared to those in set *c* (Figure 5B). Notably, when we sub-sampled set *c* to match the predicted efficacies of set *b*, the post-transcriptional regulation for set *b* greatly exceeded that of their TargetScanscore-matched counterparts from set *c* (Figure 5C). Thus, CARP enables partitioning of predicted targets with equivalent sites into true miRNA direct targets and likely downstream indirect targets. These results suggest that CARP offers substantial improvement over existing approaches. We do not rule out the possibility that some of set *c* genes could be true direct targets that CARP is missing because of failing significance threshold resulting from inherent noise in their measurements. We believe that this type of false-negative would be more prevalent for those miRNAs whose dysregulation does not lead to many indirect changes at transcription, such as miR-122, where incorporating PRO-seq would only contribute more noise to the quantification of post-transcriptional changes in the absence of substantial transcriptional change. Nonetheless, for the majority of miRNAs that we examined, we observed widespread changes in transcription (Supplementary Figure S5B), and hence it is likely that CARP does not miss many true direct targets.

In response to most miRNAs, we identified a small cohort of genes (set *f*) that experienced direct miRNA-mediated post-transcriptional down-regulation and indirect transcriptional up-regulation, resulting in minimal or no net change in mRNA abundance (Figure 5B, heatmap for set *f*). Such genes cannot be identified using transcriptome profiling alone, and such targets may represent an important and underappreciated component of miRNA biology.

Because we observed activity of ORF sites in response to miR-122 but not miR-1 (Figure 4), we next systematically examined whether this activity extended to other miRNAs. We evaluated the efficacy of ORF sites for the new set of miRNAs and compared them with the efficacy of predicted 3UTR and 5 UTR sites. We observed that different miRNAs vary substantially in their activity of ORF sites: miR-122, miR-133a, miR-155, miR-302a and miR-372 triggered post-transcriptional down-regulation of genes containing predicted ORF sites, whereas the influence of miR-1 and miR-373 on messages containing predicted ORF sites was negligible (Figure 5D). In particular, ORF sites for miR-122, miR-133a, miR-155, miR-302a and miR-372 were of comparable or greater efficacy to predicted weak 3UTR sites. Furthermore, although of lower efficacy compared to conserved 3UTR sites for all miRNAs, the conserved ORF sites for miR-122, miR-133a, miR-155 and miR-302a mediated stronger repression when compared to all predicted ORF sites, whereas miR-1, miR-372 and miR-373 lacked such a distinction (Supplementary Figure 5D). We did not observe any substantial activity of predicted 5 UTR sites for most miRNAs; the strongest evidence for effective 5UTR sites was for miR-155, although this class of sites was weaker than sites within the ORF. Taken together, these data indicate unexpected variability in the efficacy of ORF sites between different miRNAs. Although the average efficacy of these sites was substantially lower than for predicted strong 3UTR sites, for some miRNAs, we observed evidence of extensive miRNA-mediated repression via ORF sites.

Prior to our work, the conceptually equivalent approach exon-intron split analysis (EISA) (27) used pre-mRNA levels, approximated by intronic reads in RNA-seq, as a proxy for transcriptional output. To compare the performance of EISA with CARP, we contrasted changes in transcriptional output measured using PRO-seq with those inferred using intronic reads from RNA-seq. We found that changes in intronic reads in response to miR-1 were weakly correlated with changes in PRO-seq measurements (Pearson’s r between 0.11 and 0.46]; Supplementary Figure S6A). Because PRO-seq directly measures transcription by capturing transcriptionally engaged RNA polymerase (28), our results suggest that the level of intronic reads may not be a particularly accurate measure of transcription. Furthermore, we found fewer genes with significant post-transcriptional change in EISA compared to CARP for most miRNAs (Supplementary Figure S6B and C). Collectively, these findings demonstrate that CARP outperforms EISA in the quantification of post-transcriptional regulation and identification of miRNA direct targets. Nevertheless, it is clear that EISA, which requires only RNA-seq profiling, is easier to implement and is a valuable tool to define miRNA targets.

Here we demonstrate that, for a wide variety of miRNAs, CARP is able to confidently exclude a large number of false-positive predictions and selectively identify false negatives of conventional target identification approaches, in addition to robust identification of direct targets.

### Dissecting miRNA regulatory networks with CARP

Distinguishing direct miRNA targets from downstream regulatory changes is critical to gaining a systems-level understanding of miRNA gene regulatory networks. Additionally, identifying the specific direct targets (e.g., transcription factors) whose regulation results in these downstream effects is an essential component in understanding the biological roles of miRNAs. In addition to quantifying transcriptional output of genes, PRO-seq also captures and quantifies active DNA regulatory elements across the genome by measuring transcription at those elements, and has been used to identify transcription factors contributing to cell state changes by searching for transcription factor binding motifs within the differentially active elements(39). Thus, we explored whether our PRO-seq data could be used to identify transcription factors contributing to the downstream regulatory changes triggered by miRNAs.

In order to identify differential activity of DNA regulatory elements in response to miRNA induction, we first identified active elements using dREG, a tool for predicting regulatory elements using divergent transcription at active elements obtained from PRO-seq data (41). We grouped the dREG peaks into two sets: first, proximal peaks, defined as those that are close (within 1.5kb upstream and 0.5kb downstream of) to annotated transcription start sites, representing promoters; and second, the remaining, distal, peaks, representing enhancers. We found a strong correlation between the changes in transcriptional activity at dREG peaks and the changes in transcriptional output of the nearest gene, with the strongest enrichment for proximal peaks (Figure 6A and B). These results indicate that we can effectively capture changes in transcriptional activity at DNA regulatory elements which contribute to the transcriptional regulation of nearby genes.

**Figure 6.**
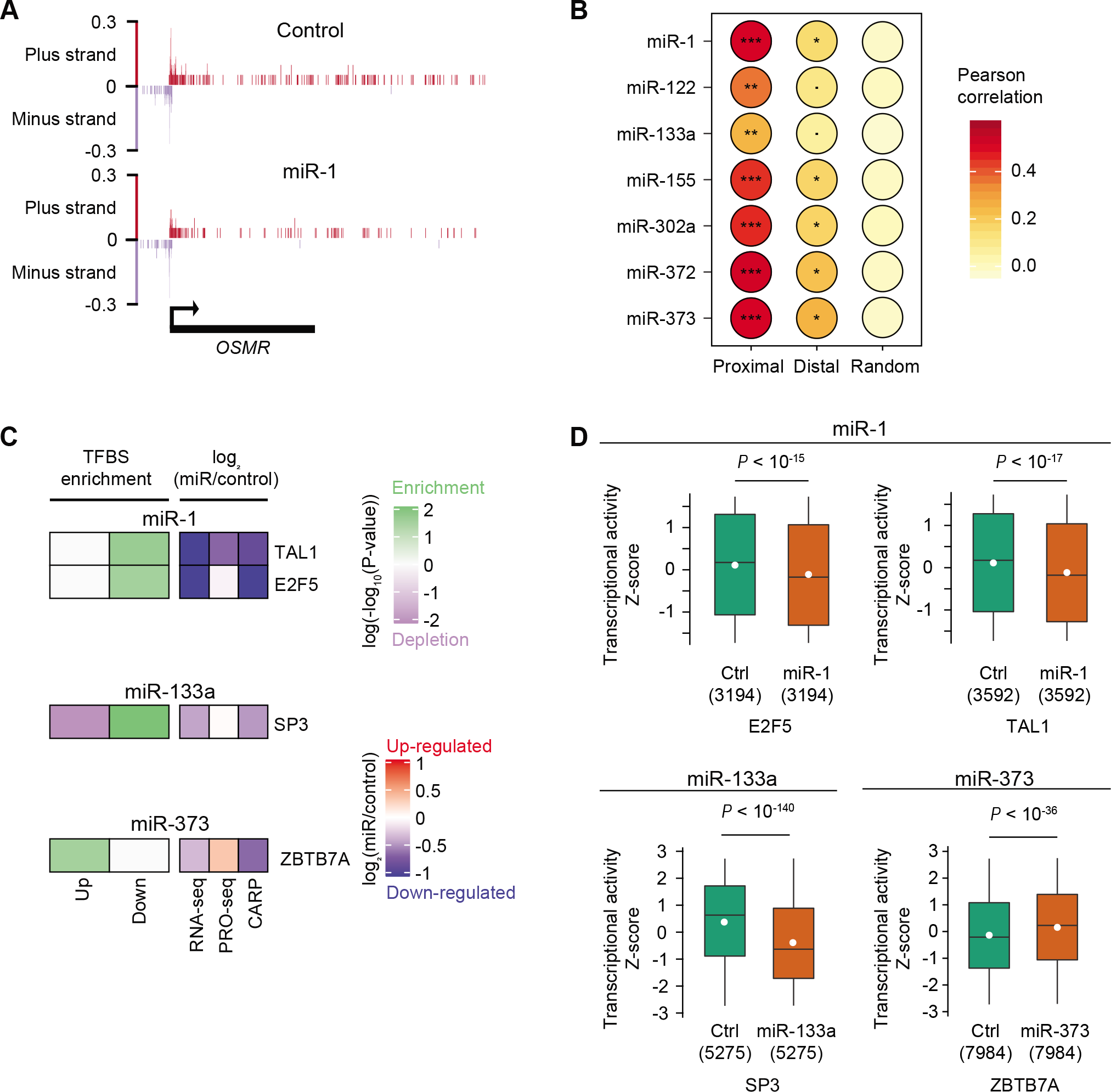
Understanding miRNA regulatory networks using CARP. (**A**) Genome browser view of PRO-seq signal (counts per million; y-axis) at the locus of the OSMR gene (bottom) demonstrating repression in promoter activity and transcriptional output in the gene-body in response to miR-1. The PRO-seq signal on plus and minus strand of the genome are depicted using red and blue colors, respectively. Divergent transcription is shown at the promoter. (**B**) Heatmap illustrating Pearson correlations between changes in transcriptional activity at proximal or distal dREG peaks with changes in transcriptional output at the nearest gene or the randomly selected gene. The correlation P values are indicated as follows: (Dot). P < 0.01, *P < 10^-10^, **P < 10^-50^, ***p < 10^-100^. (**C**) Enrichment (green) or depletion (purple) of transcription factor binding motifs in significantly up-regulated (Up) or down-regulated (Down) dREG peaks in response to respective miRNAs. Transcription factors that are direct targets of respective miRNAs (repressed in CARP with q-value < 0.05 and containing predicted 3’UTR sites) and whose binding motif was significantly enriched or depleted (FDR < 10^-3^) are depicted. For the included transcription factors, their log_2_(miR/Control) in RNA-seq, PRO-seq or CARP are color-coded using blue (down-regulated) and red (up-regulated) colors. (**D**) Distributions of relative (z-scored) transcriptional activity at putative binding sites of respective transcription factors in control cells and cells expressing miR-1 (top), miR-133a (bottom-left) or miR-373 (bottomright). The P values for indicated comparisons and the numbers of dREG peaks containing putative binding sites (x-axis; parentheses) are mentioned.

The ability to profile differential transcriptional activity of DNA regulatory elements using PRO-seq provides a unique opportunity to identify critical transcription factors embedded in the miRNA regulatory network. We hypothesized that a handful of specific transcription factors would be targeted by the cognate miRNA, and those transcription factors would be responsible for the observed downstream differences in transcription. To investigate this hypothesis, we gathered the subset of dREG peaks exhibiting significantly increased or decreased transcriptional activity and looked for enrichment of transcription factor binding motifs within these peaks. In cells expressing miR-1, we found two candidate transcription factors, TAL1 and E2F5, that are direct targets of miR-1 and whose putative binding sites were enriched in down-regulated peaks, whereas none were enriched in up-regulated peaks (Figure 6C). To more comprehensively assess these transcription factors, we evaluated their genome-wide influence on gene expression. We first examined all dREG peaks containing putative binding sites for a given transcription factor and compared their activity in cells expressing miRNAs to control cells. We found that the regulatory effects of these candidate transcription factors are widespread; for example, transcriptional activity at TAL1 binding sites was down-regulated genome-wide in cells expressing miR-1 (Figure 6D). These results suggest that miR-1 down-regulates the mRNAs of these genes directly which in turn reduces the activity of the encoded transcription factors, leading to widespread downstream transcriptional effects. Consistent with our results, previous studies have suggested that miR-1 incorporates E2F5 in its regulatory network to control cell proliferation (76,77). Similarly, we found one transcription factor, SP3, among the direct targets of miR-133a whose putative binding sites were enriched in down-regulated peaks (Figure 6C). As expected, we observed genome-wide decrease in transcriptional activity at the putative binding sites of this transcription factor (Figure 6D). We did not find any candidate transcription factors in cells expressing miR-122, consistent with our observations that there are limited indirect transcriptional effects for miR-122 (Supplementary Figure S1H). While we also did not find any candidate transcription factors for miR-155, miR-302a or miR-372, likely because there were very few dREG peaks with significant change in activity, we found one candidate transcription factor, ZBTB7A, among the direct targets of miR-373, whose putative binding sites were enriched in those dREG peaks that exhibited increased transcriptional activity (Figure 6C). ZBTB7A has been shown to acts as a transcriptional repressor (78), indicating that miR-373-mediated post-transcriptional repression of ZBTB7A promotes transcriptional up-regulation of its target genes. Consistent with this interpretation, we observed genome-wide activation of transcriptional activity at the putative binding sites for ZBTB7A (Figure 6D). We note that changes in activity at many dREG peaks was modest (Figure 6D), consistent with miRNAs acting to fine-tune levels of targets. Taken together, we have shown that PRO-seq not only measures transcriptional outputs, but also captures differentially active DNA regulatory elements, allowing us to perform motif enrichment analysis and thereby identify candidate transcription factors responsible for genome-wide indirect effects of miRNAs. We believe that this additional benefit of PRO-seq will be of substantial value for gaining a comprehensive understanding of the impact of miRNAs at a systems-level.

## DISCUSSION

This study was motivated by the assumption that many miRNAs with consequential functions work, in part, by eliciting substantial downstream regulatory changes, and that most such changes would occur via transcriptional regulation. That is, miRNAs likely function, to some extent, by controlling one or more transcription factors. Our data, and many published studies, corroborate this assumption. Identifying the direct targets of a miRNA and distinguishing these direct targets from downstream changes (indirect targets) is challenging. These challenges derive from (i) the subtle nature of miRNA regulation, (ii) false positives and false negatives in target prediction, (iii) the large number of potential targets, and (iv) the potential for an extensive number of downstream indirect targets. Inevitably, some downstream targets will possess putative target sites, and will often be erroneously considered as direct targets. Our work highlights the extent of such errors, together with many other aspects of miRNA-controlled regulatory networks that are difficult to parse without directly measuring transcriptional regulation that is triggered by a miRNA. The combination of miRNA target prediction and RNA-seq profiling is routinely applied to the study of miRNAs. This study demonstrates that adding PRO-seq (or related tools that measure transcription across the genome) provides far more reliable definition of the target set of a miRNA, with significant added insights into the overall regulatory network controlled by a miRNA.

An important aspect of this study is the reliance on cell lines that ectopically express miRNAs within the physiological range, rather than using miRNA transfection, as is common in earlier work examining the regulatory impact of miRNAs. Indeed, perhaps the extent of false positives that exist amongst predicted targets derives, in part, from the dependence on training datasets that used transiently transfected miRNAs that likely exceed physiological levels. Another important aspect of this study is the use of multiple genomic tools to profile the regulatory changes mediated by a miRNA – namely, RNA-seq, PRO-seq, ribosome profiling and AGO eCLIP-seq, generating a comprehensive dataset of orthogonal approaches that encapsulates almost all aspects of miRNA-mediated targeting and regulation. Finally, we have provided combined RNA-seq and PRO-seq data for five additional miRNAs; in total, these data serve as an ideal resource to aid in our understanding of miRNA target sites and in further improvement of prediction algorithms.

The precision with which targets are identified by CARP allows us to more reliably examine non-canonical miRNA targeting; that is, targeting that extends beyond seed-type sites within 3’UTRs. In particular, we have found that certain miRNAs (e.g., miR-122) have large numbers of functional target sites within coding sequence, whose efficacy is comparable to many canonical 3’UTR sites. Importantly, we validated this observation using AGO eCLIP-seq. In contrast, the suite of target sites for other miRNAs including miR-1 and miR-373, were restricted to their conventional location within 3’UTRs. We note that other studies have also hinted at miRNA-specific differences in targeting propensities (15). Although there is certainly precedence for miRNA coding sites, we are not aware of compelling evidence that certain miRNAs possess large numbers of effective sites within coding sequence, nor studies that robustly compare the extent of such targeting between a cohort of different miRNAs. Thus, two important conclusions from this study are that the targeting properties of miRNAs are not uniform, and that for some miRNAs, a substantial fraction of their regulatory impact is mediated by target sites within coding sequence. It is also worth noting that for miR-122, sites within the ORF appear to potentiate repression of genes with predicted weak 3’UTR sites, whereas there is no additional repression from ORF sites for genes with predicted strong 3’UTR sites. Future studies will be needed to decipher the mechanistic bases for these observations.

The relative contributions of translational repression and accelerated decay due to miRNA-mediated regulation is an important question, both because of mechanistic implications, and, pragmatically, due to the reliance on transcriptome profiling in the vast majority of miRNA studies, including CARP. Here, we have investigated two miRNAs, miR-1 and miR-122, using a suite of tools that provides an ideal dataset to rigorously quantify the relative contributions of decay and translational regulation elicited by miRNAs. We estimate that less than 15% of total regulation occurs via translational regulation; moreover, targets that are regulated exclusively via translational repression are exceedingly rare. Therefore, CARP is well-suited for the identification of true miRNA targets, except for the rare contexts such as the Zebrafish embryo (64,65) where miRNA direct targets are subjected exclusively to translational repression without any change in mRNA stability.

Discovery of indirect targets is equally important for the understanding of miRNA-mediated regulation as the knowledge of miRNA direct targets. We discovered substantial indirect regulation at transcription for many miRNAs; for example, expression of miR-1 triggered transcriptional up-regulation of 133 genes and transcriptional down-regulation of 237 genes. Interestingly, not all of the indirect targeting we observed in such cases was transcriptional; we also found a set of genes that are indirectly regulated at the post-transcriptional level. It is important to note that direct regulation by miRNAs is unlikely to account for this class of targets, because we saw no increase in relative AGO density in mRNA of genes subject to post-transcriptional regulation that lack canonical target sites, and, for miR-1, many of these post-transcriptionally changing genes were up-regulated. For miR-1, the extent of this regulation was surprisingly widespread; while we identified 370 genes changing at transcription, we estimate 273 genes to be indirectly post-transcriptionally regulated. This indicates that miRNAs are likely embedded in complex regulatory networks comprised of both transcriptional and post-transcriptional regulation.

A major advantage of CARP is its ability to simultaneously measure transcriptional and post-transcriptional changes. One set of genes that illustrates this importance is the set of high confidence direct targets, across multiple miRNAs, which show little or no change in mRNA abundance. Our data indicate that these targets are simultaneously directly repressed and indirectly activated. Identifying such targets previously was challenging. Overall, such targets constitute 2% of the genes subject to miRNA-mediated post-transcriptional regulation. We argue that the study of these miRNA targets is important in order to understand how miRNAs regulate a complete gene regulatory network. Although this study was not intended to investigate the biological roles of any of the miRNAs we used, the frequency with which we found miRNA targets whose repression is balanced by transcriptional up-regulation suggests that such targets may also be common in native regulatory pathways involving miRNAs.

The primary motivation for this study was to demonstrate the utility of PRO-seq, in combination with RNA-seq, to robustly identify post-transcriptional regulation, and thereby robustly identify miRNA targets. However, in addition to genome-wide quantification of transcription, an integral aspect of PRO-seq data is genome-wide quantification of promoter and enhancer activities. This additional feature greatly amplifies the utility of PRO-seq in understanding the regulatory impact of a miRNA. Presumably, many gene regulatory pathways incorporate both miRNAs and transcription factors, and the combination of RNA-seq with PRO-seq is clearly well justified in effectively separating regulatory contributions of miRNAs from those mediated by transcription factors. This combination identifies genes subject to transcriptional and post-transcriptional control, and provides a profile of active DNA regulatory elements genome-wide. The ability of CARP to provide this information makes it an ideal approach for probing complex regulatory networks. Taken together, we show that CARP provides a framework for simultaneously measuring regulation occurring at multiple stages of gene expression, which could prove to be a powerful approach for comprehensively disentangling miRNA gene regulatory networks *in vivo*.

## DATA AVAILABILITY

The RNA-seq, PRO-seq, ribosome profiling, eCLIP-seq and small RNA-seq data reported in this paper are available from GEO (GSE140367).

## ACKNOWLEDGEMENTS

We thank Jennifer K. Grenier and the Transcriptional Regulation and Expression Facility (TREx) for helpful discussion and assistance with RNA-seq library generation, Rene Geissler for technical assistance with eCLIP and ribosome profiling protocols, members of the Grimson laboratory for helpful discussions, and John Lis and his laboratory for helpful discussions regarding PRO-seq protocols.

## Author Contributions

R.P. and A.G. conceived this project. R.P. performed most of the experiments and all of the analyses. J.W. performed eCLIP and ribosome profiling experiments, Y.J. assisted R.P. with PRO-seq experiments. E.F. performed luciferase reporter assays. R.P. and A.G. wrote the paper with assistance from J.W.

## Graphical Abstract

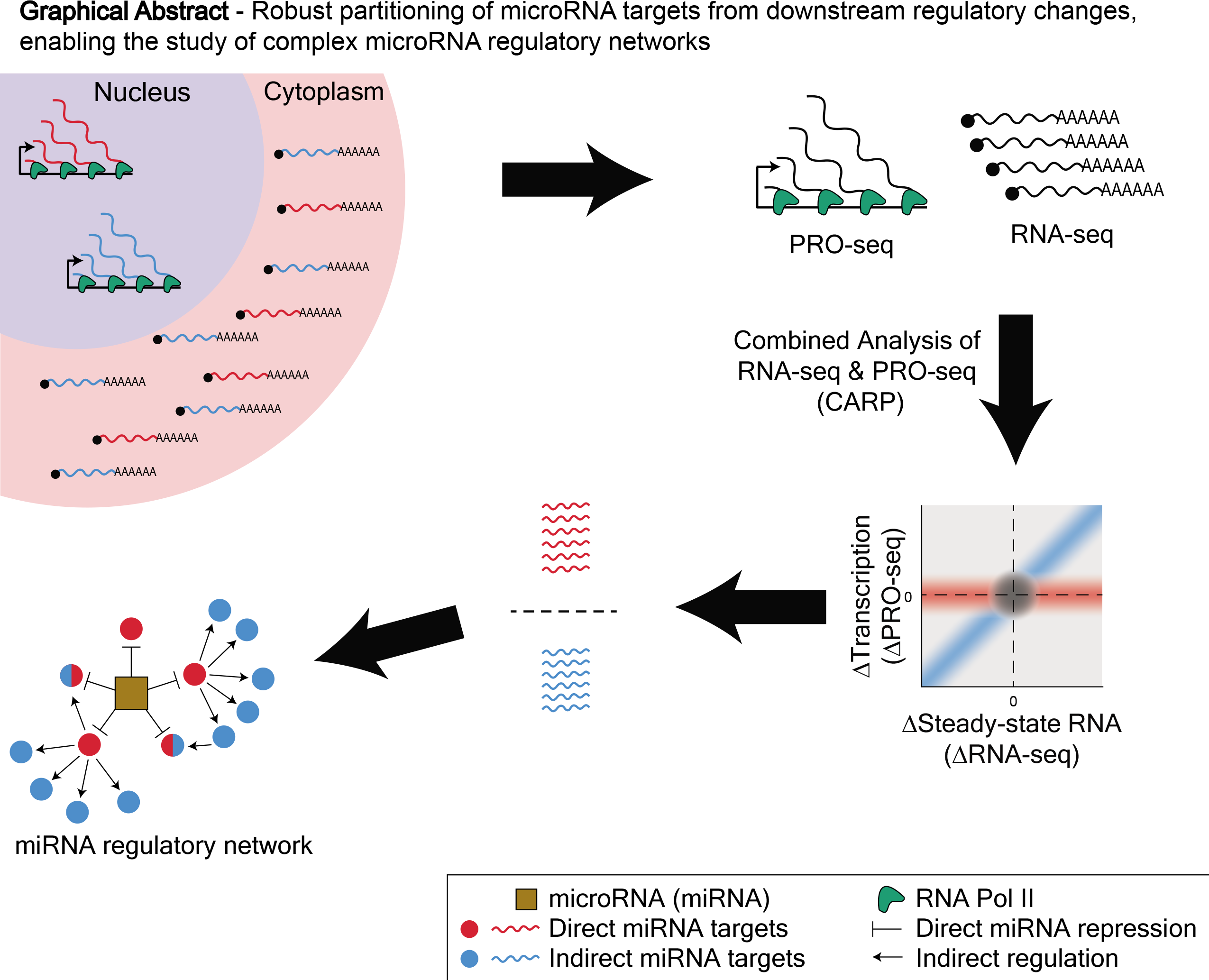
Robust partitioning of microRNA targets from downstream regulatory changes, enabling the study of complex microRNA regulatory networks

