## Supplementary Data for "Robust partitioning of microRNA targets from downstream regulatory changes"

### Supplementary Figure S1

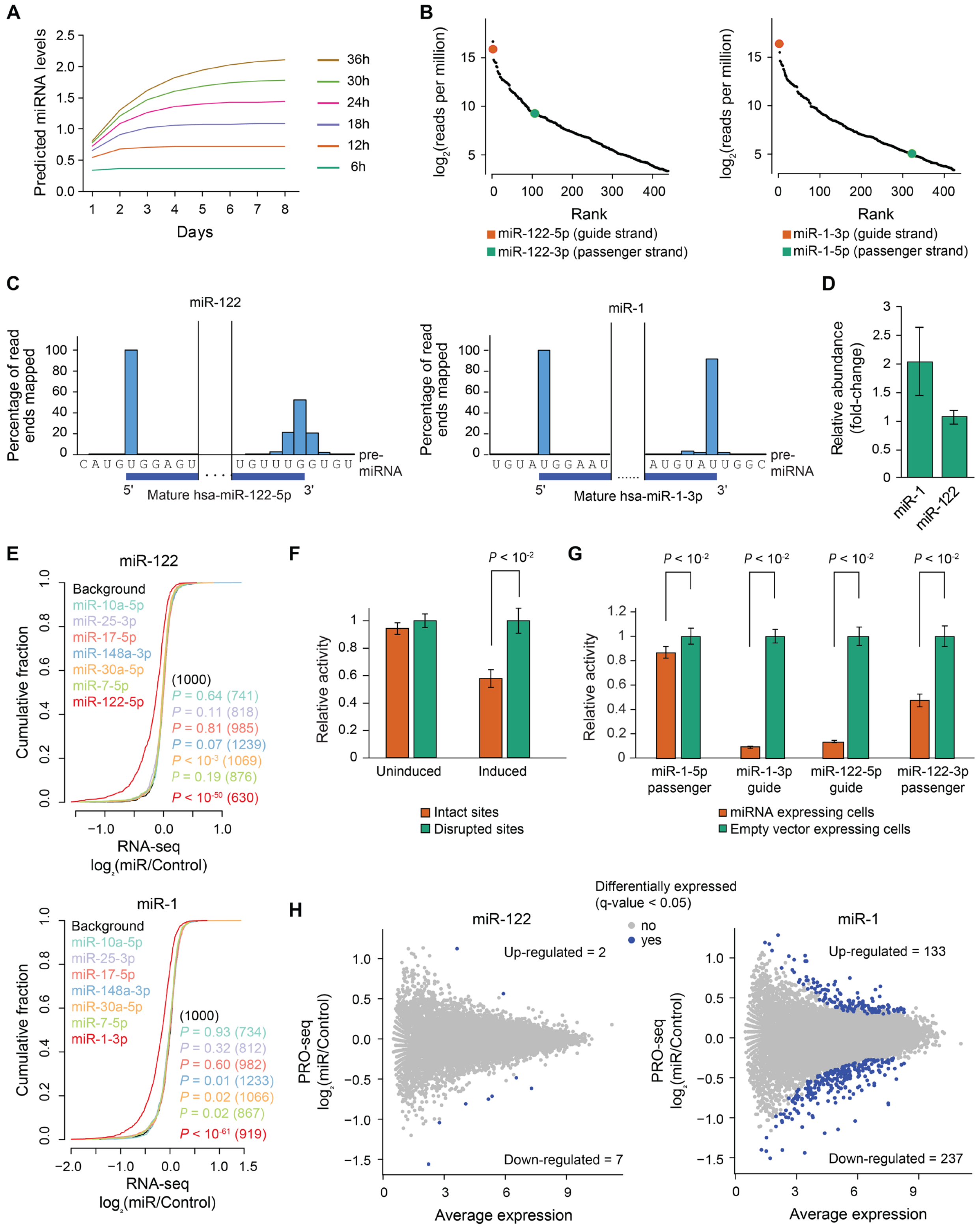

**Supplementary Figure S1. Analysis of expression level and processing of induced miRNAs.** (**A**) Line graph depicting modeling of steady-state levels (x-axis) for miRNAs with different half-lives (color-coded). (**B**) Expression profiling of miRNAs in cells expressing miR-122 (left) or miR-1 (right) using small RNA-seq. (**C**) Bar plots representing relative levels of alternative isoforms of the mature guide strands of miR-122 (left) and miR-1 (right). The percentages of 5′ end (left of each chart) and 3′ end (right of each chart) of small RNA-seq reads mapping to the pre-miRNA are plotted. The sequence of the pre-miRNA is shown below each bar, with the 5′ and 3′ ends of the mature strand depicted by the blue bar. (**D**) qPCR quantification of relative levels of induced miRNAs compared to the most highly detected endogenous miRNA, miR-10. Absolute levels of each miRNA were determined using a standard curve. The error bars represent standard deviation across six replicates. (**E**) Cumulative distribution plots of RNA-seq changes for predicted strong targets (TargetScan context++ score < -0.2) of endogenous miRNAs in response to miR-122 (top) or miR-1 (bottom). The predicted strong targets of highly-detected endogenous miRNAs (small RNA-seq) and the cognate miRNA were compared with 1000 randomly selected genes (background). The P value and number of genes in each set (parentheses) are indicated by color. (**F**) Luciferase reporter assays to monitor induced miRNA efficacy. The reporter construct contains two miR-1 3′UTR target sites, or mutant target sites, and was assayed in cells with and without miR-1 induction. The errors bars represent standard deviation across six replicates. (**G**) Luciferase reporter assays to evaluate loading of the correct miRNA strand (guide strand). The reporter constructs contained a target site perfectly matched to the complete sequence of either the guide or passenger strand of miR-1 or miR-122, and were assayed in cells with or without the expression of cognate miRNA. The errors bars represent standard deviation across six replicates. (**H**) MA plots representing change in transcriptional output (measured with PRO-seq) upon induction of miR-122 (left) or miR-1 (right) compared to control. Significantly changing genes are colored in blue. The number of up-regulated and down-regulated genes are indicated.

### Supplementary Figure S2

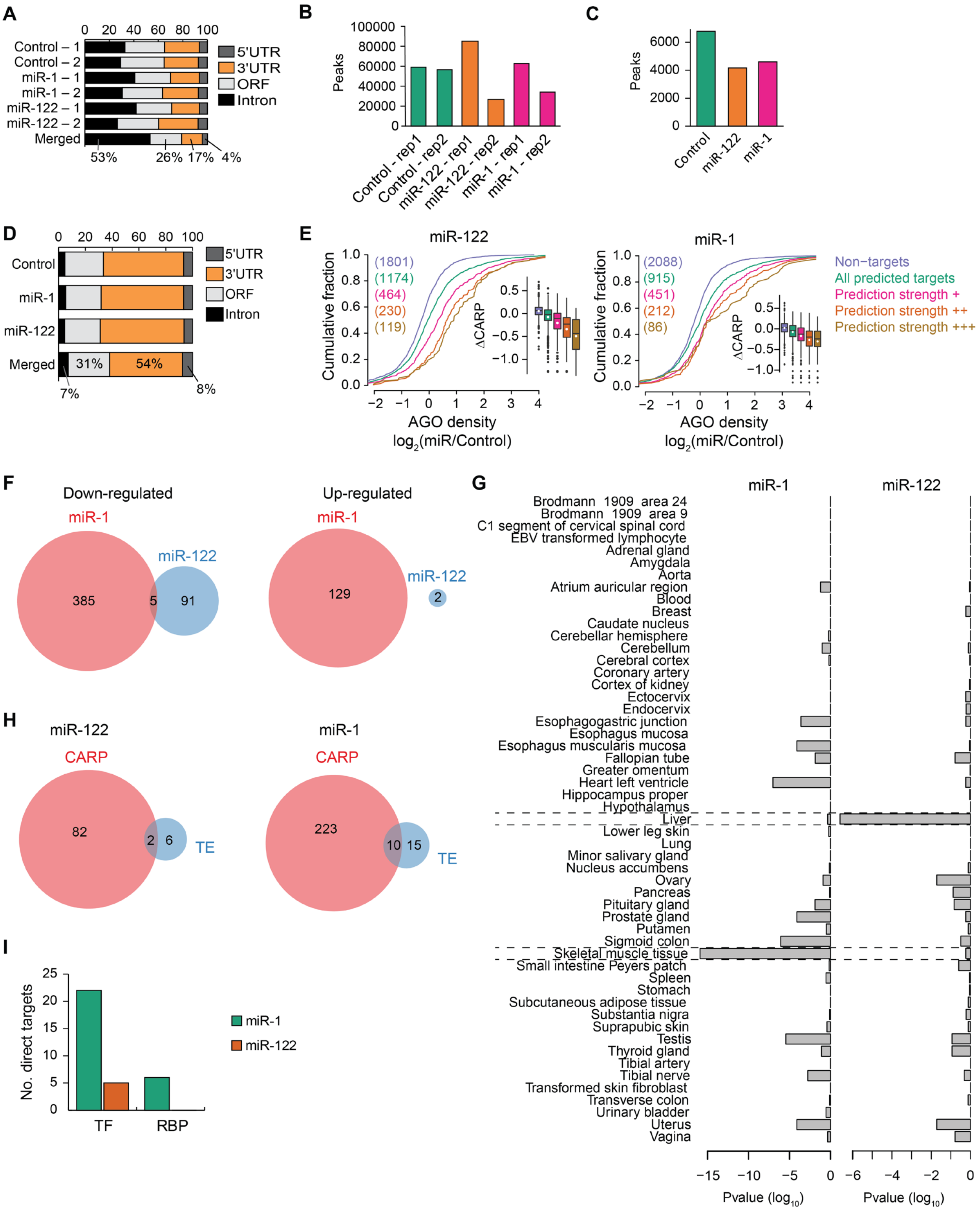

**Supplementary Figure S2. Quality filtering of AGO eCLIP data and efficacy of CARP and EISA.** (**A**) Stacked bar plots depicting fractions of AGO eCLIP peaks in the 5′UTR, 3′UTR, ORF and intron for cells expressing miR-1, miR-122, or the empty vector control. (**B**) Bar plots demonstrating variability in number of eCLIP peaks between replicates. (**C**) Number of eCLIP peaks after removing irreproducible peaks using irreproducibility discovery rate (IDR) analysis. (**D**) Stacked bar plots representing fractions of AGO eCLIP peaks in the 5′UTR, 3′UTR, ORF and intron for cells expressing miR-1, miR-122, or the empty vector control, after performing IDR analysis. (**E**) Cumulative distribution plots depicting increase in relative AGO density $\log_2$(miR/Control) at eCLIP peaks in cells expressing miR-122 (left) or miR-1 (right) with increasing prediction strength of targets. The insets represent post-transcriptional regulation of targets ($\Delta$CARP, y-axis) with increasing prediction strength of targets. The prediction strengths +, ++ and +++ correspond to sets targets with context++ score < -0.1, < -0.2 and < -0.3, respectively (**F**) Venn diagrams depicting overlap of genes post-transcriptionally down-regulated (left) or up-regulated (right) in response to miR-1 and miR-122. (**G**) Tissue-specificity analysis of CARP-identified direct targets. Bar plots represent P values for testing whether the direct targets of miR-1 (left) or miR-122 (right) are down-regulated compared to 5,000 randomly selected genes in each of the 53 tissues from GTEx (y-axis). The P values are calculated using one-sided Wilcoxon rank sum test and corrected using the Benjamini-Hochberg method. (**H**) Venn diagrams comparing number of predicted targets (including strong and weak) exhibiting significant repression either at the level of mRNA stability (measured using CARP) or at translation (measured using ribosome profiling). (**I**) Number of direct targets of miR-1 or miR-122 that code for putative transcription factors (TF; as defined in CIS-BP (1)) or RNA binding proteins (RBP; as defined in ATtRACT (2)).

### Supplementary Figure S3

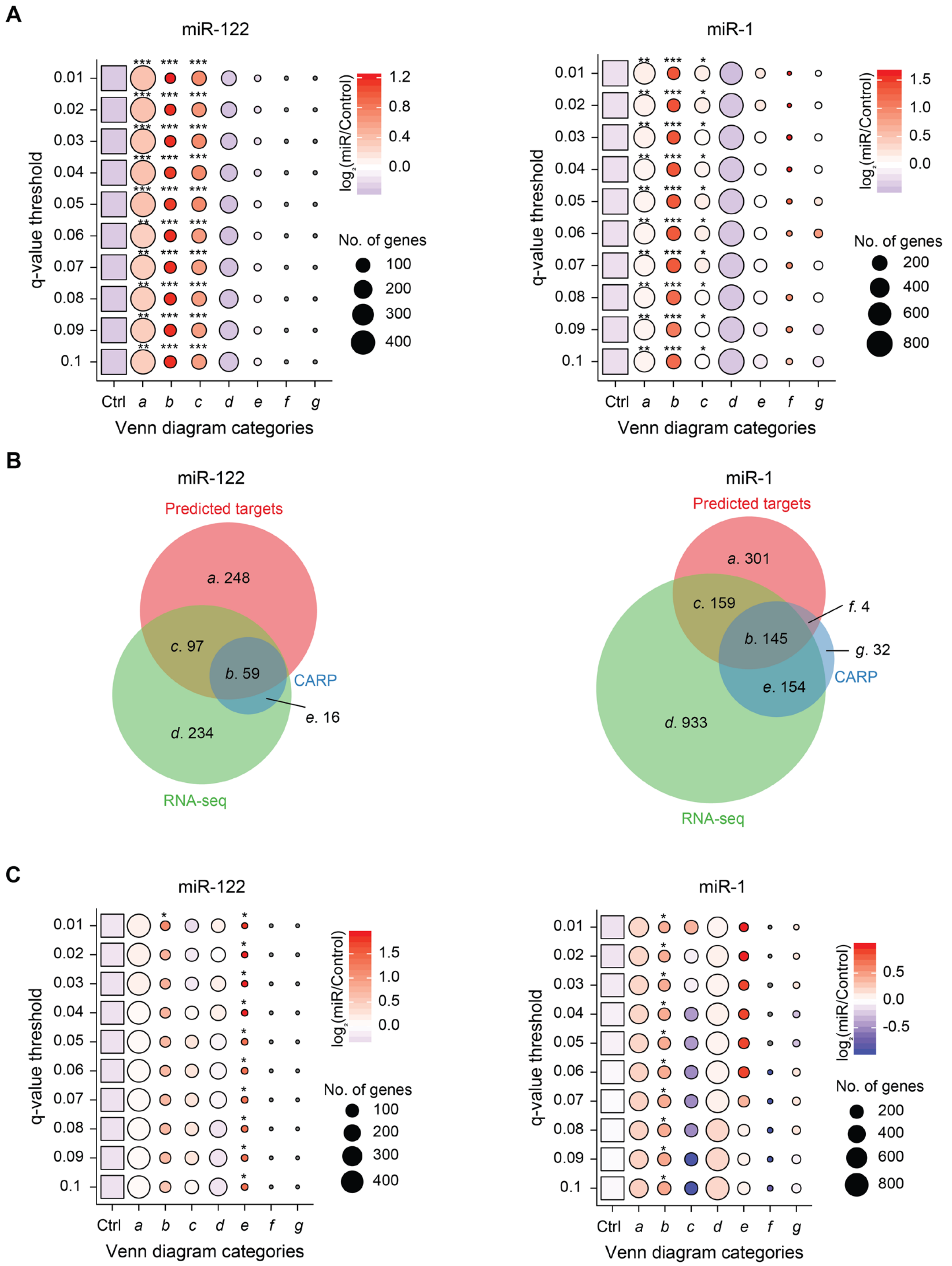

**Supplementary Figure S3. Robustness of CARP across different significance thresholds and alternative polyadenylation status.** (**A**) Relative AGO density and number of significantly post-transcriptionally repressed genes across a range of q-value thresholds for miR-122 (left) and miR-1 (right) for each Venn diagram category *a*-*g* (as described in Figure 3A). The blue-red color gradient represents median of relative AGO density ($\log_2$(miR/Control)) at eCLIP peaks within 3′UTRs, and the size of each bubble depicts number of genes within each gene-set. The relative AGO density for every gene-set (bubble) for a given q-value threshold was compared to the relative AGO density for the corresponding control set (square); the control set is as described in Figure 4B, but for different q-value thresholds. P values are indicated as follows: *P $< 0.01$, **P $< 10^{-5}$, ***P $< 10^{-10}$. (**B**) Venn diagrams are identical to those in Figure 3A, except that the predicted strong targets that lost predicted target sites due to alternative cleavage and polyadenylation (predicted using GETUTR (3)) are excluded. (**C**) Same as in A, but representing relative AGO density at eCLIP peaks in the ORF.

### Supplementary Figure S4

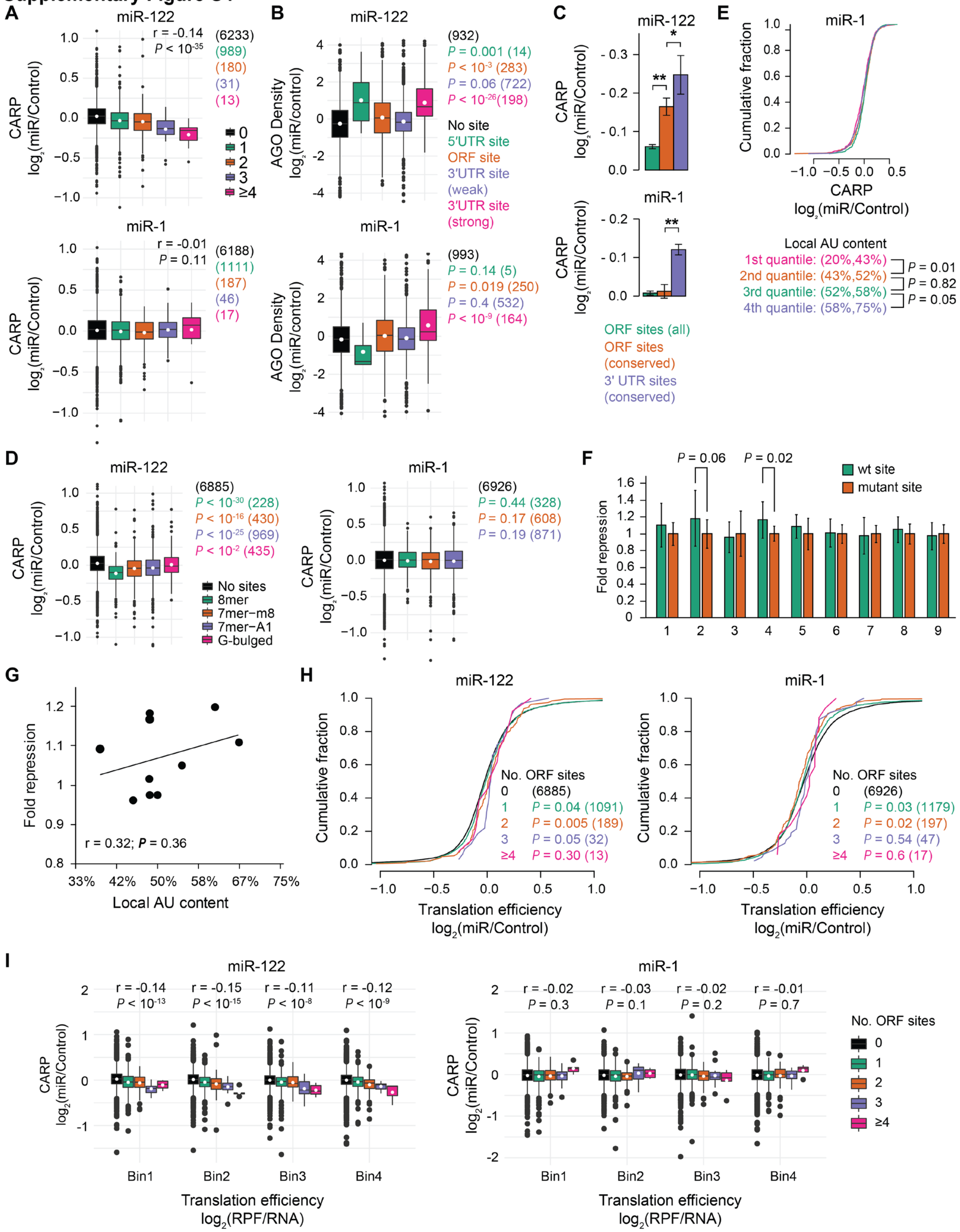

**Supplementary Figure S4. Evaluation of ORF site efficacy.** (**A**) Boxplots depicting change in post-transcriptional regulation (CARP, y-axis) for genes with increasing number of predicted ORF sites to miR-122 (top) or miR-1 (bottom). Genes that contain predicted 3′UTR sites are excluded. Number of genes in each set are indicated in parentheses. Pearson correlation coefficients (r) and P values (P) between number of predicted ORF sites and change in post-transcriptional regulation are indicted. (**B**) Relative AGO density (miR/Control) at eCLIP peaks in different regions of an mRNA. Each set contains genes with predicted target sites (matches to 8mer, 7mer-m8 or 7mer-A1 site motifs) exclusively in that region. Each set was compared to the set of genes containing no sites. The P values and number of genes in each set (parentheses) are indicated. (**C**) Bar plots depicting median change in post-transcriptional regulation of genes in response to miR-122 (top) or miR-1 (bottom). The activities of genes containing predicted sites in ORF (as described in Figure 4A), predicted conserved sites in ORF (PACCMIT-CDS: $P\_SH < 0.05$) or predicted conserved sites in 3′UTR (TargetScan: $Pct > 0.3$) were normalized by the median change observed for genes without any predicted target sites. The error bars represent standard error. P values are indicated as follows: $*P < 0.05$, $**P < 10^{-4}$. (**D**) Post-transcriptional changes mediated by different types of predicted target sites within the ORF. Genes that contain predicted 3′UTR sites are excluded. Each set was compared to the set of genes containing no sites. The P value and number of genes in each set (parentheses) are indicated. (**E**) Relationship between post-transcriptional regulation and local AU content around predicted miR-1 ORF sites, as described in Figure 4B. (**F**) Luciferase reporter assays to assess the efficacy of additional candidate miR-122 ORF sites (1-9), plotted as described in Figure 4C (n=12). (**G**) Scatter plot indicating relationship between local AU content (as described in Figure 4B) and fold repression from Figure 4C and Supplementary Figure S4F. A linear fit to the data (black line), Pearson correlation coefficient (r) and the associated P value are indicated. (**H**) Cumulative distribution plots of translational efficiency of genes containing increasing numbers of predicted ORF sites for miR-122 (left) or miR-1 (right). Genes that contain predicted 3′UTR sites are excluded. (**I**) Evaluation of ORF site efficacy for genes with various levels of translation. Genes were grouped into four equal size bins according to the translational efficiency (calculated by normalizing ribosome protected fragments (ribosome profiling) with mRNA levels (RNA-seq)) and their post-transcriptional regulation (CARP; y-axis) is plotted as a function of different numbers of predicted ORF sites. Pearson correlation coefficients (r) and P values (P) between number of predicted ORF sites and change in post-transcriptional regulation are indicated.

**Supplementary Figure S5**

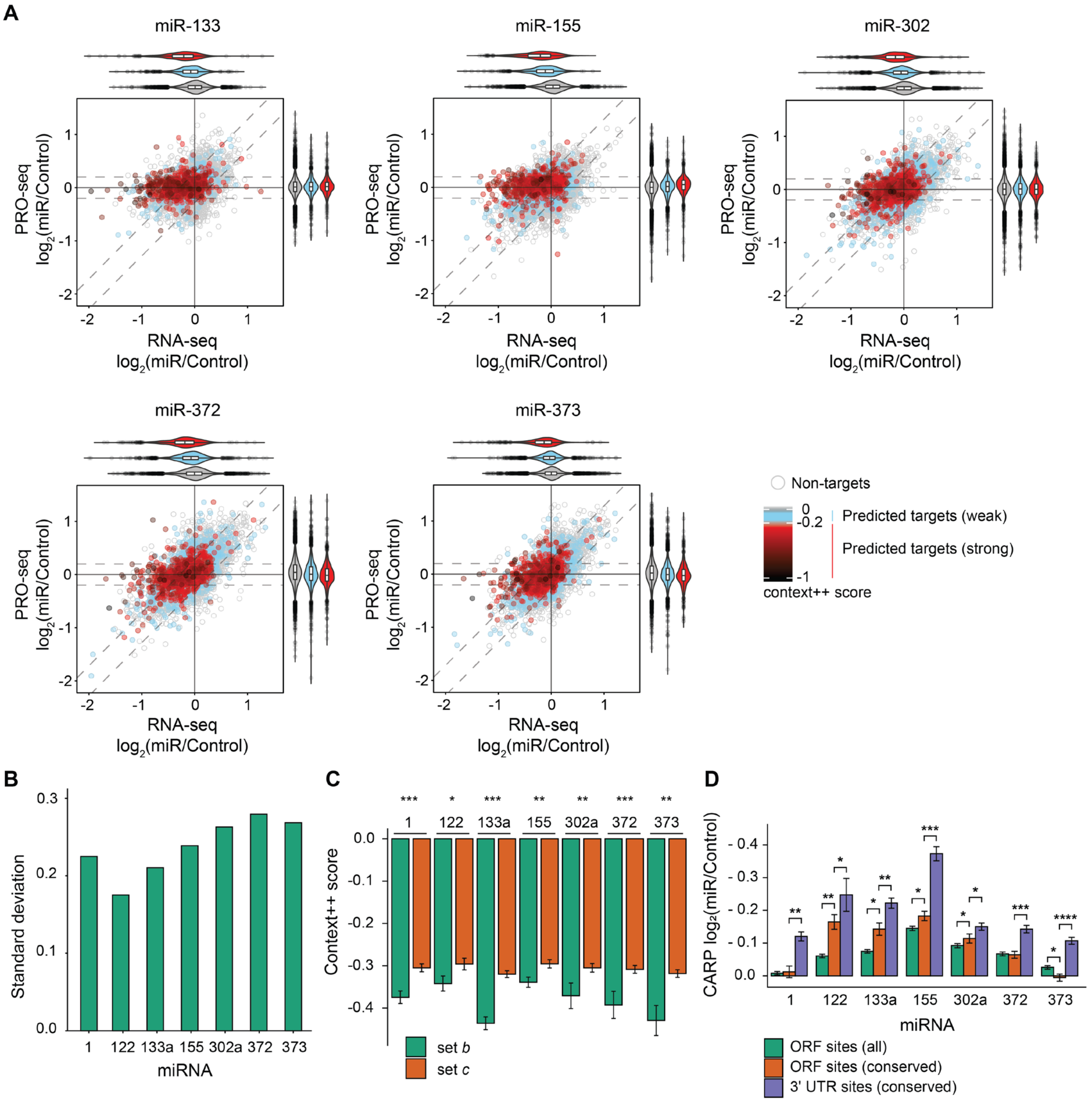


**Supplementary Figure S5. Post-transcriptional and transcriptional regulation mediated by different miRNAs used in this study.** (**A**) Dot plots depicting changes in mRNA abundance and transcriptional output for miR-133a, miR-155, miR-302a, miR-372 and miR-373, otherwise as described in Figure 1C. (**B**) Degree of transcriptional regulation as measured using standard deviation of changes in transcriptional output in response to specific miRNAs (PRO-seq $\log_2$(miR/Control)) across all genes. (**C**) Median TargetScan prediction efficacy (context++ score) for set *b* and set *c* genes; these sets correspond to the Venn diagram categories depicted in

Figure 5B. P values are indicated as follows: *P < 0.05, **P < 0.005, ***P < 0.0005. (**D**) Bar plots depicting median change in post-transcriptional regulation of genes in response to different miRNAs, same as described in panel S4A. P values are indicated as follows: *P < 0.05, **P < $10^{-4}$, ***P < $10^{-8}$, ****P < $10^{-12}$.

### Supplementary Figure S6

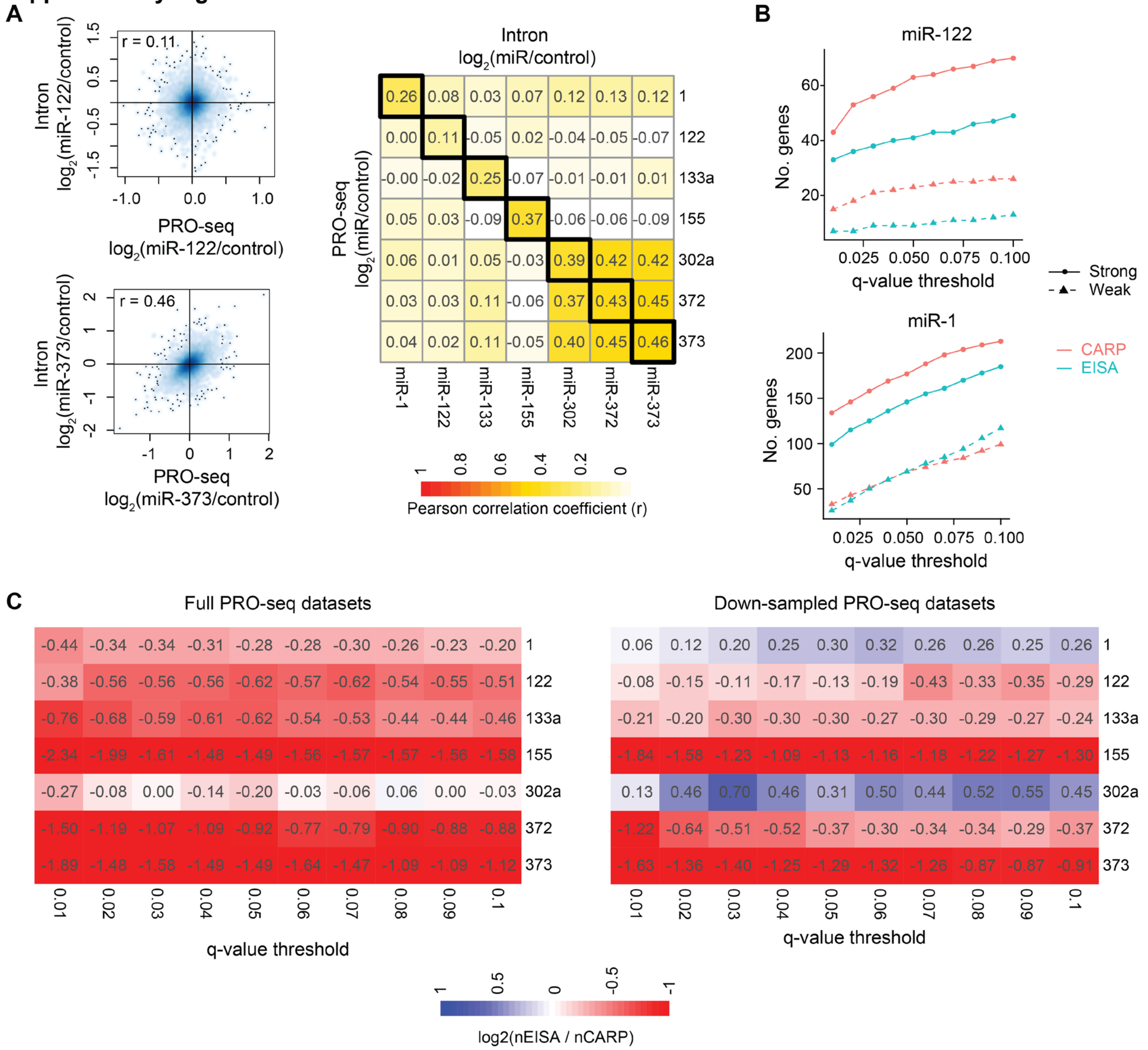


**Supplementary Figure S6. Comparison between efficacy of CARP and Exon-intron split analysis.** (**A**) Scatter plots comparing change in transcriptional output as measured using PRO-seq (x-axis) or as inferred using intronic reads (y-axis) in response to miR-122 (top) or miR-373 (bottom). Dark blue to light blue color gradient depicts high to low density of data points. The points represent outliers of the density estimation model. Pearson correlation coefficients (r) are indicated for each comparison. Heatmap (right) summarizes Pearson correlation coefficients for all possible combinations of miRNAs used in this study. The diagonal includes correlations for same miRNAs between the two methods. (**B**) The number of predicted targets (strong and weak) that are significantly repressed at post-transcription in cells expressing miR-122 (top) or miR-1 (bottom) compared to control cells for a range of significance (q-value) thresholds according to CARP (red) or EISA (blue). (**C**) Heatmaps summarizing the differences between CARP and EISA (as in B) for predicted strong targets of all miRNAs used in this study either using the full PRO-seq datasets (left) or the down-sampled PRO-seq data to match the sequencing depth of intron data (right). The color key represents ratio ($log_2$) of number of predicted strong targets that are identified as post-transcriptionally repressed using EISA (nEISA) and the number of those identified using CARP (nCARP).

**Supplementary Table S1.** Sequences of miRNA hairpins that were introduced in miRNA-expression cassette, with the XbaI site at the 5' end and the XmaI site at the 3' end. The backbone of the hairpins is adopted from one of the hairpins ("A5") used in (4)(Fang,Bartel,etal-2015). The underscored nucleotides indicate expected mature miRNA 5p and 3p sequences.

>Hsa-miR-1
TCTAGACCTTTGCTGTTCTCCAAAACATACTTCTTTATATGCCCATAGTGTCTTAATTGTATGG
AATGTAAAGAAGTATGTATTGGCGAACGGCTTTATCGTCGGTCCCGGG

>Hsa-miR-122
TCTAGACCTTTGCTGTTCTCCATGTGGAGTGTGACAATGGTGTTTGGTGTCTTAATTGCAAACG
CCATTATCACACTAAATATGGCGAACGGCTTTATCGTCGGTCCCGGG

>Hsa-miR-133a
TCTAGACCTTTGCTGTTCTCCACAGCTGGTAAAATGGAACCAAATCGTGTCTTAATTGGATTTG
GTCCCCTTCAACCAGCTGTGGCGAACGGCTTTATCGTCGGTCCCGGG

>Hsa-miR-155
TCTAGACCTTTGCTGTTCTCCATGTTAATGCTAATCGTGATAGGGGTTGTGTCTTAATTGGACT
CCTACATATTAGCATTAACATGGCGAACGGCTTTATCGTCGGTCCCGGG

>Hsa-miR-302a
TCTAGACCTTTGCTGTTCTCCACCACTTAAACGTGGATGTACTTGCTGTGTCTTAATTGAGTAA
GTGCTTCCATGTTTTGGTGATGGCGAACGGCTTTATCGTCGGTCCCGGG

>Hsa-miR-372
TCTAGACCTTTGCTGTTCTCCAGGCCTCAAATGTGGAGCACTATTCTGTGTCTTAATTGAGAAA
GTGCTGCGACATTTGAGCGTCTGGCGAACGGCTTTATCGTCGGTCCCGGG

>Hsa-miR-373
TCTAGACCTTTGCTGTTCTCCAATACTCAAAATGGGGGCGCTTTCCGTGTCTTAATTGGGGAAG
TGCTTCGATTTTGGGGTGTTGGCGAACGGCTTTATCGTCGGTCCCGGG

**Supplementary Table S2.** Oligos and primers used for qPCR (Supplementary Figure S1D).
RT primer:
`Uni. RT primer 5'CAGGTCCAGTTTTTTTTTTTTTTTVN`

Universal reverse primer:
`Uni. miRPCR_rv 5'CAGGTCCAGTTTTTTTTTTTTTTT`

MiRNA-specific forward primers:
```
miR1-3p_fwWTail      5'GCAGTGGAATGTAAAGAAGTA
miR122-5p_fwWTail    5'ACAGTGGAGTGTGACAATG
miR10-5p_fwWTail     5'AGTACCCTGTAGATCCGAA
Inert2.2-5p_fwWTail 5'CAGTAAAAATCGCGTGGAT
```

Synthetic templates for standard curve:
```
miR1-3p_annotEnd_synTmplt
     5'CAGGTCCAGTTTTTTTTTTTTTTTATACATACTTCTTTACATTCCA
miR122-5p_annotEnd_synTmplt
     5'CAGGTCCAGTTTTTTTTTTTTTTTCAAACACCATTGTCACACTCCA
miR10-5p_annotEnd_synTmplt
     5'CAGGTCCAGTTTTTTTTTTTTTTTCACAAATTCGGATCTACAGGGTA
Inert2.2-5p_annotEnd_synTmplt
     5'CAGGTCCAGTTTTTTTTTTTTTTTCATTAATCCACGCGATTTTTA
```

Spike-in RNA oligo:
`Inert2.2-5p    5'UAAAAAUCGCGUGGAUUAAUG`

**Supplementary Table S3.** DNA sequence of 3'UTR fragments cloned downstream of luciferase for assays in Supplementary Figure S1F. The SacI site is at the 5' end and the XbaI site at the 3' end. The underscored nucleotides represent miR-1 sites.

>miR-1 WT sites: GLCCI1 SacI-XbaI [pAG76]

```
GAGCTCTATTTTATGCATTTCCCTTTCCTCATTACATTCCACATTCTTAGAATAAGAAGTGCAT
TCAATCCTAGGAGAATGATAATCCTGGACATGGGTGAACATGAGGAGAACCAGCAAAATCTGTG
GTGTTTGACATCACTTTGTCATGTGGTTACAAGTAAAACAACTGTTGCATTCACTGTTTCAACA
TGTGTACATGTGGCTTTTTTAAAAGTTCAGGTGTTGCTCAGTAAAGGACTGTGACAATGTTGCA
AATAAAGTGTTCAGTACTGGACTGTACATAAACATTCCACATTGTGTCTAGA
```

>miR-1 disrupted sites: GLCCI1 SacI-XbaI [pAG77]

```
GAGCTCTATTTTATGCATTTCCCTTTCCTCATTAGAATGCACATTCTTAGAATAAGAAGTGCAT
TCAATCCTAGGAGAATGATAATCCTGGACATGGGTGAACATGAGGAGAACCAGCAAAATCTGTG
GTGTTTGACATCACTTTGTCATGTGGTTACAAGTAAAACAACTGTTGCATTCACTGTTTCAACA
TGTGTACATGTGGCTTTTTTAAAAGTTCAGGTGTTGCTCAGTAAAGGACTGTGACAATGTTGCA
AATAAAGTGTTCAGTACTGGACTGTACATAAAGAATGCACATTGTGTCTAGA
```

**Supplementary Table S4.** DNA sequences cloned downstream of luciferase, used as siRNA-type (fully complementary) target sites for assays in Supplementary Figure S1G. The XbaI site is at the 5' end and the SalI site at the 3' end.

>hsa-miR-1-1_5p
TCTAGATATGGGCATATAAAGAAGTATGTGTCGAC

>hsa-miR-1-1_3p
TCTAGAATACATACTTCTTTACATTCCAGTCGAC

>hsa-miR-122_5p
TCTAGACAAACACCATTGTCACACTCCAGTCGAC

>hsa-miR-122_3p
TCTAGATATTTAGTGTGATAATGGCGTTGTCGAC

**Supplementary Table S5.** Luciferase reporter sequences for miR-122 ORF site assays in Figure 4C and Supplementary Figure S4F. Shown below is the firefly luciferase ORF sequence (from pmirGLO) with the inserted linker sequence (encoding for amino acids GGGSGGGS) underlined. The asterisk denotes where the endogenous coding regions containing miR-122 ORF sites were inserted. The inserted sequences (for both the wildtype and mutant target site) are listed below, along with gene names.

>pmirGLO_fireflyLucCDS_withLinker
ATGGAAGATGCCAAAAACATTAAGAAGGGCCCAGCGCCATTCTACCCACTCGAAGACGGGACCG
CCGGCGAGCAGCTGCACAAAGCCATGAAGCGCTACGCCCTGGTGCCCGGCACCATCGCCTTTAC
CGACGCACATATCGAGGTGGACATTACCTACGCCGAGTACTTCGAGATGAGCGTTCGGCTGGCA
GAAGCTATGAAGCGCTATGGGCTGAATACAAACCATCGGATCGTGGTGTGCAGCGAGAATAGCT
TGCAGTTCTTCATGCCCGTGTTGGGTGCCCTGTTCATCGGTGTGGCTGTGGCCCCAGCTAACGA
CATCTACAACGAGCGCGAGCTGCTGAACAGCATGGGCATCAGCCAGCCCACCGTCGTATTCGTG
AGCAAGAAAGGGCTGCAAAAGATCCTCAACGTGCAAAAGAAGCTACCGATCATACAAAAGATCA
TCATCATGGATAGCAAGACCGACTACCAGGGCTTCCAAAGCATGTACACCTTCGTGACTTCCCA
TTTGCCACCCGGCTTCAACGAGTACGACTTCGTGCCCGAGAGCTTCGACCGGGACAAAACCATC
GCCCTGATCATGAACAGTAGTGGCAGTACCGGATTGCCCAAGGGCGTAGCCCTACCGCACCGCA
CCGCTTGTGTCCGATTCAGTCATGCCCGCGACCCCATCTTCGGCAACCAGATCATCCCCGACAC
CGCTATCCTCAGCGTGGTGCCATTTCACCACGGCTTCGGCATGTTCACCACGCTGGGCTACTTG
ATCTGCGGCTTTCGGGTCGTGCTCATGTACCGCTTCGAGGAGGAGCTATTCTTGCGCAGCTTGC
AAGACTATAAGATTCAATCTGCCCTGCTGGTGCCCACACTATTTAGCTTCTTCGCTAAGAGCAC
TCTCATCGACAAGTACGACCTAAGCAACTTGCACGAGATCGCCAGCGGCGGGGCGCCGCTCAGC
AAGGAGGTAGGTGAGGCCGTGGCCAAACGCTTCCACCTACCAGGCATCCGCCAGGGCTACGGCC
TGACAGAAACAACCAGCGCCATTCTGATCACCCCCGAAGGGGACGACAAGCCTGGCGCAGTAGG
CAAGGTGGTGCCCTTCTTCGAGGCTAAGGTGGTGGACTTGGACACCGGTAAGACACTGGGTGTG
AACCAGCGCGGCGAGCTGTGCGTCCGTGGCCCCATGATCATGAGCGGCTACGTTAACAACCCCG
AGGCTACAAACGCTCTCATCGACAAGGACGGCTGGCTGCACAGCGGCGACATCGCCTACTGGGA
CGAGGACGAGCACTTCTTCATCGTGGACCGGCTGAAGAGCCTGATCAAATACAAGGGCTACCAG
GTAGCCCCAGCCGAACTGGAGAGCATCCTGCTGCAACACCCCAACATCTTCGACGCCGGGGTCG
CCGGCCTGCCCGACGACGATGCCGGCGAGCTGCCCGCCGCAGTCGTCGTGCTGGAACACGGTAA
AACCATGACCGAGAAGGAGATCGTGGACTATGTGGCCAGCCAGGTTACAACCGCCAAGAAGCTG
CGCGGTGGTGTTGTGTTCGTGGACGAGGTGCCTAAAGGACTGACCGGCAAGTTGGACGCCCGCA
AGATCCGCGAGATTCTCATTAAGGCCAAGAAGGGCGGCAAGATCGCCGTGGGCGGAGGATCTGG
AGGCGGATCT*TAA

>site1_ SAMD15_wt; target site is underlined
AGTGAAGAATCAATTGGTACACATTATGAGTTTTTGCAACCACTCCAAAAATTGCTTAATGTCA
GTGAAGAATGCTCA

>site1_ SAMD15_mut; target site is underlined
AGTGAAGAATCAATTGGTACACATTATGAGTTTTTGCAACCTCTGCAAAAATTGCTTAATGTCA
GTGAAGAATGCTCA

>site2_BACH1_wt; target site is underlined
CACCTAGCAAAAGGCTTCTGGAGTGACATTTGCAGCACGGACACTCCTTGCCAAATGCAGTTAT
CACCTGCTGTGGCC

>site2_BACH1_mut; target site is underlined
CACCTAGCAAAAGGCTTCTGGAGTGACATTTGCAGCACGGATACACCTTGCCAAATGCAGTTAT
CACCTGCTGTGGCC

>site3_CUX1_wt; target site is underlined
AGAAAGCGGCTTATCGAACAGAGCCGGGAGTTCAAGAAGAACACTCCAGAGGATTTGCGCAAGC
AGGTAGCGCCGCTG

>site3_CUX1_mut; target site is underlined
AGAAAGCGGCTTATCGAACAGAGCCGGGAGTTCAAGAAGAATACACCTGAGGATTTGCGCAAGC
AGGTAGCGCCGCTG

>site4_CARD6_wt; target site is underlined
TCTCAGTTCAAATCCGATCAGTCCAACCCATCCACAGTCAAACACTCCCAGCCTAAACCCTTCC
ATTCTGTGCCCTCT

>site4_CARD6_mut; target site is underlined
TCTCAGTTCAAATCCGATCAGTCCAACCCATCCACAGTCAAGCATAGCCAGCCTAAACCCTTCC
ATTCTGTGCCCTCT

>site5_DCHS1_wt; target site is underlined
AAGGCCTTCCGCATCCACCCCCAGACTGGAGAAGTGACCACACTCCAAACCCTGGACCGTGAGC
AGCAGAGCAGCTAT

>site5_DCHS1_mut; target site is underlined
AAGGCCTTCCGCATCCACCCCCAGACTGGAGAAGTGACCACTCTGCAAACCCTGGACCGTGAGC
AGCAGAGCAGCTAT

>site6_CYP24A1_wt; target site is underlined
GCCATCAAAACAATGATGAGCACGTTTGGGAGGATGATGGTCACTCCAGTCGAGCTGCACAAGA
GCCTCAACACCAAG

>site6_CYP24A1_mut; target site is underlined
GCCATCAAAACAATGATGAGCACGTTTGGGAGGATGATGGTCACACCTGTCGAGCTGCACAAGA
GCCTCAACACCAAG

>site7_CHGA_wt; target site is underlined
TCCAAGCCCAGCCCCATGCCTGTCAGCCAGGAATGTTTTGAGACACTCCGAGGAGATGAACGGA
TCCTTTCCATTCTG

>site7_CHGA _mut; target site is underlined
TCCAAGCCCAGCCCCATGCCTGTCAGCCAGGAATGTTTTGAGACTCTGCGAGGAGATGAACGGA
TCCTTTCCATTCTG

>site8_TWISTNB_wt; target site is underlined

GTGGACAGTGGTACCACAAAGCTAGCAGATGATGCAGATGACACTCCAATGGAAGAGTCAGCCC
TGCAGAATACTAAT

>site8_TWISTNB _mut; target site is underlined
GTGGACAGTGGTACCACAAAGCTAGCAGATGATGCAGATGATACACCTATGGAAGAGTCAGCCC
TGCAGAATACTAAT

>site9_SLC4A8_wt; target site is underlined
GGAGAGTTCATGGGATCTGCGTGCGGCCATCATGGACCCTACACTCCTGATGTCCTCTTTTGGT
CCTGTATTCTCTTT

>site9_SLC4A8_mut; target site is underlined
GGAGAGTTCATGGGATCTGCGTGCGGCCATCATGGACCCTATACACCTGATGTCCTCTTTTGGT
CCTGTATTCTCTTT

>site10_RWDD2A_wt; Figure 4C; target site is underlined
ACAAGGGAGGCGCTGCCACCAAAAATCGAATTTGTAATTACACTCCAGATTGAAGAGCCCAAGG
TGAAAATTGATTTG

>site10_RWDD2A _mut; Figure 4C; target site is underlined
ACAAGGGAGGCGCTGCCACCAAAAATCGAATTTGTAATTACTCTGCAGATTGAAGAGCCCAAGG
TGAAAATTGATTTG

**Supplementary Table S6.** Synthetic miRNA duplexes used for verifying miR-122 ORF sites in luciferase assays (Figure 4C and Supplementary Figure S4F).

inert2.2 (control miRNA mimic)

```
  5'-UAAAAAUCGCGUGGAUUAAUG-3'
     ||||||.|||||..|||||
3'-UUAUUUUUGGCGCAUUUAAUU-5'
```

hsa-miR-122

```
  5'-UGGAGUGUGACAAUGGUGUUUG-3'
     |  ||||||| |||||||||
3'-AUAAAUCACACUAUUACCACAA-5'
```
